# The Unreasonable Effectiveness of Cell Types in Describing Neuronal Physiological Features

**DOI:** 10.64898/2026.08.27.744753

**Authors:** Harshil Sharma, Xiao-Ping Liu, Thomas Chartrand, Brian Kalmbach, Ed Lein, Stefan Mihalas, Zhixin Lu

**Author notes:** These authors contributed equally to this work (Stefan Mihalas, Zhixin Lu).

## Abstract

Single-cell RNA sequencing (scRNA-seq) captures detailed gene expression profiles at scale, while patch-clamp recordings measure intrinsic neuronal electrophysiological properties. Modeling the relations between these two modalities remains a challenge. Here, we compare how well electrophysiological features can be predicted by traditional transcriptomic cell type classification, representations derived from a foundational model (scGPT) pretrained on large-scale scRNA-seq datasets, ion channel-coding genes, and highly variable genes. Using paired transcriptomic and electrophysiological patch-sequencing data from 495 human neurons from neurosurgical tissue, we find that cluster-level cell type representations generally outperform highly variable gene selection, ion channel gene selection, and context-enriched scGPT embeddings. Notably, performance varies across model architectures and initializations, and the best results are obtained by combining the outputs of separate cell type and scGPT-based models. Together, these findings suggest that traditional discrete cellular classification is highly effective in predicting physiological features. For maximum performance it can be complemented by pretrained transformer models.

**Author summary:** Understanding how a neuron’s genes relate to its electrical properties is a major goal in neuroscience. New technologies now make it possible to measure gene expression in individual cells and, simultaneously, to record how those same cells respond to electrical signals. However, relating these two types of information at the single cell level remains difficult. In this study, we tested whether a modern artificial intelligence model trained on large collections of gene expression data could help connect gene activity to electrical behavior in human brain cells. We compared this approach with simpler strategies, such as using selected sets of genes or grouping cells by their known types. We found that basic cell type descriptions often predicted electrical properties better than more complex gene-based methods alone. The strongest results were achieved by combining cell type knowledge with information from the artificial intelligence model and training them together. These findings suggest that when linking data is in the hundreds, simpler representations outperform large general-purpose AI models in isolation, but the two approaches may be complementary rather than competing.

## Introduction

Single-cell RNA sequencing (scRNA-seq) has enabled large-scale profiling of neuronal gene expression across diverse brain regions and cell types, generating datasets comprising tens of thousands to millions of cells. In contrast, electrophysiological measurements—which directly characterize neuronal functional properties such as excitability and firing dynamics—remain low-throughput and experimentally demanding. Consequently, the ability to accurately predict electrophysiological parameters from scRNA-seq data would greatly enhance the functional interpretability of existing transcriptomic atlases and provide a powerful avenue for studying neuronal diversity at scale.

A unique opportunity to investigate this problem is provided by patch-sequencing (Patch-seq), a multimodal experimental approach that combines patch-clamp electrophysiology with scRNA-seq measurements in the same cell (as reviewed in [1]). Patch-seq datasets enable direct empirical assessment of relationships between gene expression and electrophysiological phenotypes. However, due to experimental complexity, Patch-seq remains limited in scale, particularly for human neurons. As a result, existing human Patch-seq datasets can be viewed as critical testbeds for evaluating and comparing different approaches to predicting electrophysiological features from scRNA-seq data. Such comparisons help clarify which *transcriptomic representations* are most informative for electrophysiological prediction and provide reference results for the development of improved models as larger datasets become available.

Predicting electrophysiological properties from transcriptomic profiles is intrinsically challenging. The relationship between gene expression and cell physiology is indirect, mediated by protein concentrations and localization, alternative splicing, accessory subunit composition, post-translational regulation, and is further obscured by technical and biological noise. Traditional approaches rely on subsets of highly variable genes or clustering algorithms such as Seurat for cell type labeling, which can only partially capture this complexity. To better contextualize this problem, prior research can be broadly categorized along two axes: model expressivity (rigid to flexible) and species (mouse versus human).

Early studies in mouse datasets used linear regression to identify replicable correlations between ion channel gene expression and specific firing properties [2–5]. Subsequent biophysical modeling of mouse visual cortex neurons demonstrated that ion channel composition inferred from electrophysiological properties aligns with the expression of corresponding genes [6]. Cadwell et al. (2016) extended the examined gene space via generalized linear models (GLMs) to partially predict electrophysiological features from transcriptomic profiles [7]. Collectively, these studies on simplified models support the existence of transcriptomic signatures underlying electrophysiological phenotypes, but remain constrained to small subsets of genes and have limited explained variance.

Moreover, these constrained models determined largely consistent cell classification between the two modalities, insofar as their small datasets granted [6, 7]. Indeed, descriptive work on larger collections of cells across multiple regions of the mouse cortex reveals that while electrophysiology-based and transcriptomic-based cell typing yield good correspondence with one another, both t-types and e-types exhibit noticeable heterogeneity, with neuronal features varying along a continuum rather than forming sharply separated types [8–13]. These findings underscore the need for more flexible, nonparametric modeling approaches capable of capturing continuous structure across modalities.

Motivated by this insight, recent work has applied nonlinear deep learning architectures to cross-modal prediction. Both coupled autoencoders and variational autoencoders (VAEs) have been used to learn shared latent representations between transcriptomic and electrophysiological data, improving cross-modal alignment [14, 15]. Complementary work has demonstrated the inverse mapping: transcriptomic cell type can be recovered directly from electrophysiological time series using deep neural networks [16]. Together, these studies establish that expressive models can leverage subtle features within each modality to recover information about the other.

Nevertheless, these modeling efforts are limited to murine datasets, and human modeling remains underexplored. Human and mouse cortical neurons show extensive transcriptomic and physiological divergence, even among homologous cell types, suggesting limited cross-species generalization [17, 18]. This limitation is consequential because human-specific gene-to-physiology models could help interpret disease-associated transcriptional changes in terms of altered neuronal function, including in Alzheimer’s disease and autism spectrum disorder, where sequencing studies have revealed cell-type-specific molecular alterations [19, 20]. While Bardy et al. (2016) applied both linear and nonlinear models to neurons derived from human induced pluripotent stem cells (iPSCs) to identify gene–electrophysiology relationships, this work was constrained by developmental immaturity and limited physiological diversity of iPSC-derived cells [21].

The gap between the biological complexity of human neurons and the limited scope of existing models remains striking—especially in light of recent advances in transformer-based foundation models for single-cell biology, capable of learning transferable, multimodal cellular representations (see [22, 23] for comprehensive reviews of this topic). scGPT is one such transformer-based foundation model pretrained on large scale human scRNA data. It represents each cell as a sequence of gene tokens, where each token combines learned embeddings of gene identity and discretized expression values, optionally augmented with meta-information embeddings. A special classification token aggregates gene-level information into a cell-level representation. During its self-supervised generative pretraining, scGPT iteratively decodes masked gene expression values from the input in an autoregressive manner. The final classification token output from this process serves as the cell embedding—a 512-dimensional vector that encapsulates the learned transcriptomic representation of the cell. These cell embeddings proved useful for downstream tasks such as cell type annotation, and when combined with gene-level embeddings, captured aspects of the gene regulatory network. When fine-tuned on additional data, scGPT was able to predict outcomes of gene perturbation and integrate additional data modalities [24]. However, it remains unclear whether these cell embeddings can capture functional cellular properties beyond transcriptional identity, such as electrophysiological behavior.

In this study, we present a comparative benchmark of transcriptomic representations for predicting electrophysiological parameters from human Patch-seq data. Using a dataset of 495 human neurons with paired snRNA-seq and electrophysiological recordings, we trained various multilayer perceptron (MLP) models to predict electrophysiological features from the following transcriptomic representations: (i) scGPT-derived cell embeddings, expression levels of the top 512 highly variable genes, (iii) expression of ion channel-coding genes, and (iv) cell type information encoded as one-hot vectors. We further evaluated combinations of gene-based representations with cell type information.

Rather than proposing a new predictive model, our goal is, with the limited data, to provide a systematic and controlled comparison of these representations under a unified experimental framework. By quantifying their relative predictive performance on the same human dataset, we aim to clarify the strengths and limitations of each approach and to establish reference results that can guide future studies. As larger and more diverse human Patch-seq datasets become available, such benchmarks will be essential for informing the development of improved models linking transcriptomic variation to neuronal function.

## Materials and methods

### Ethics statement

This study is a secondary analysis of previously published, de-identified human data; no new human tissue or data were collected. Neurosurgical tissue was originally obtained with informed consent under protocols approved by the institutional review boards of the participating hospitals (Seattle - Harborview Medical Center, Swedish Medical Center, and University of Washington Medical Center; Amsterdam - Vrije Universiteit Medical Center; and Szeged - University of Szeged), as described in the original publications [18, 25].

### Data used

Data for this study were obtained from Patch-seq characterizations of human neocortical layer 1 (L1) GABAergic interneurons and layer 2/3 (L2/3) glutamatergic supragranular neurons [18, 25], collected by the Allen Institute for Brain Science. These datasets include matched electrophysiological, transcriptomic, and metadata annotations for each recorded neuron. Although these datasets were generated using single-nucleus RNA-seq (snRNA-seq) rather than scRNA-seq, prior studies indicate that the two methods yield broadly concordant transcriptomic profiles of human samples [26–28].

Cell type assignments for Patch-seq samples were adopted from the original publications and are briefly summarized here [18, 25]. For the L2/3 dataset, the Seurat v3 pipeline was used to integrate Patch-seq transcriptomic profiles with the reference snRNA-seq dataset and perform label transfer [29]. For the L1 dataset, Patch-seq samples were mapped to the reference taxonomy using a hierarchical, tree-based method. At each branch point, correlations between the Patch-seq sample and the marker genes defining each branch were calculated, and the branch with the highest correlation was selected. The process was repeated iteratively until reaching a terminal leaf of the dendrogram, yielding the final cell type call. After combining the L1 and L2/3 datasets, cell types represented by fewer than four cells were excluded, resulting in a final dataset of 495 neurons spanning 15 cell types.

Electrophysiological features were calculated from recordings in the original publications, a subset of which is illustrated in Fig 1 [18, 25, 30]. After combining the two datasets and retaining only electrophysiological features common to both, 15 shared features were available. All action potential features and one additional feature, latency, were excluded from subsequent analyses due to consistently poor predictability. Latency was also excluded because of its idiosyncratic nature, leaving eight electrophysiological features for this study, listed in Table 1. These features were percentile-normalized across cells to mitigate differences in scale.

**Table 1.** Table describing the electrophysiological features used in the study.

| Feature Name | Units | Description |
| --- | --- | --- |
| sag ratio | – | Difference between the minimum membrane potential after hyperpolarization and the steady-state potential, divided by the peak deflection during the stimulus |
| resting membrane potential | mV | Average of the pre-stimulus membrane potential. |
| rheobase | pA | Minimum current amplitude of one-second-long steps that evoked an action potential. |
| f-I slope | Hz/pA | Slope of linear fit to the frequency response of the cell versus stimulus intensity curve. |
| input resistance | MΩ | Resistance of cell membrane as measured by a linear fit to responses to hyperpolarizing current steps. |
| time constant $\tau$ | sec | Time constant of exponential fit to responses to hyperpolarizing current steps. |
| mean firing rate | Hz | Average firing rate across the entire stimulus interval. |
| adaptation | – | Rate at which firing speeds up or slows down during a stimulus. |

**Fig 1.**
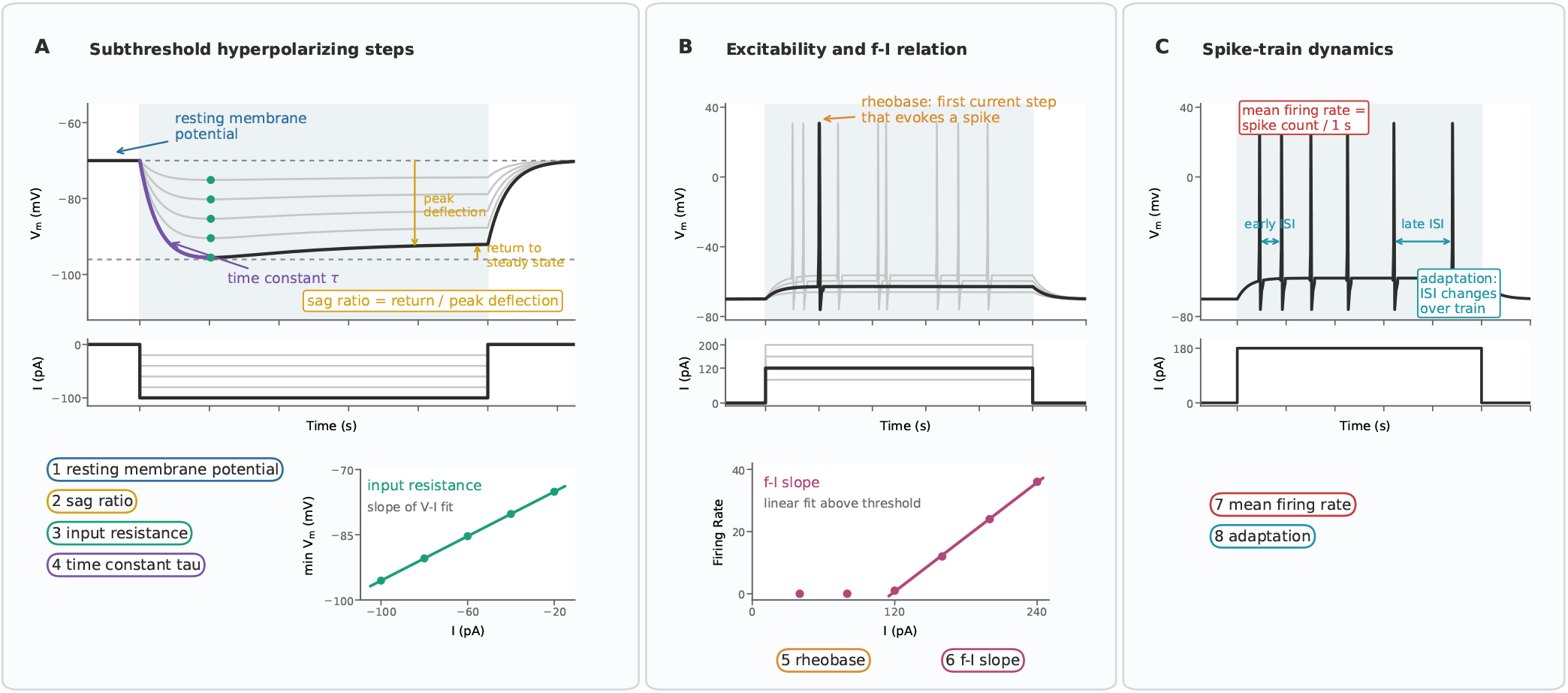
Electrophysiological features extracted from current-clamp recordings. (A) Hyperpolarizing long-square current steps were used to estimate resting membrane potential, sag ratio, input resistance, and membrane time constant. Resting membrane potential was measured from the pre-stimulus baseline; sag ratio was defined as the return from the minimum membrane potential to steady state divided by the peak deflection; input resistance was estimated from the slope of the voltage-current relation; and the membrane time constant was obtained from exponential fits to subthreshold voltage responses. (B) Depolarizing long-square steps were used to identify rheobase, defined as the smallest current step that evoked an action potential, and to compute the f-I slope from the suprathreshold firing-rate-current relation. (C) Representative suprathreshold responses were used to compute mean firing rate from spike count over the stimulus interval and adaptation from changes in interspike intervals across the spike train.

Transcriptomic data initially contained log-transformed combined intron and exon read counts for 50,281 genes per cell. However, these were filtered to contain only 30,682 genes, overlapping with the vocabulary of scGPT. Among these 30,682, we identified the top 512 highly variable genes using Scanpy and 328 ion channel-coding genes using the list provided by the HGNC database [31, 32]. Nine genes were shared between the highly variable and ion channel gene sets. A breakdown of the transcriptomic data used is given in Table 2.

**Table 2.** Breakdown of cells studied and genes used in creating each transcriptomic representation.

| Metric | Value |
| --- | --- |
| Total genes from SMART-Seq | 50,281 |
| Genes used in scGPT | 30,682 |
| scGPT embedding dimensionality | 512 |
| Highly variable genes | 512 |
| ion channel-coding genes | 328 |
| Total cells | 495 |
| Layer 1 cells | 229 |
| Layer 2/3 cells | 266 |
| Cell types identified by Seurat clustering | 15 |

We opted to use the scGPT model pretrained on 13.2 million human brain cells, which we found slightly outperformed the whole human model pretrained on 33 million cells spanning the whole body (see S16 Fig). For each cell, this pretrained model ingested the full transcriptome (the 30,682 genes) and produced a 512-dimensional cell embedding [24].

Units of measurement are provided for reference only. All values were percentile-normalized for our analysis, and raw measurement units were not directly predicted.

Summary of the genes used to construct each transcriptomic representation (top) and of the cells included in the study (bottom). Transcriptomic data was obtained using SMART-Seq v4, providing intron+exon counts for 50,281 genes. These were filtered to the 30,682 genes present in the scGPT vocabulary, all of which were used to generate 512-dimensional cell embeddings. For comparison with the scGPT embeddings, electrophysiological features were also predicted directly from the top 512 highly variable genes (HVGs), 328 ion channel-coding genes (9 of which overlapped with the HVGs), and 15 cell types identified by Seurat clustering. In total, 495 neurons with both transcriptomic data and eight electrophysiological features were included in the study.

### Models and training

These data provide four distinct representations of cells and their potential relationship to electrophysiology. Two representations are established standards based in highly variable genes and cell types, one is a biologically motivated representation focused on ion channel-coding genes, and the fourth is a novel representation derived from a pretrained foundation model. To enable a simple and direct comparison of their ability to predict electrophysiological properties, we initially trained separate multilayer perceptron (MLP) models, each using a single representation as input. In later tests, representations were combined with cell type information.

Formally, for each cell *i*, the model takes as input a representation *x*_*i*_ ∈ ℝ^*d*^ and produces a prediction *ŷ*_*i*_ ∈ ℝ^8^. The MLP with *L* hidden layers is defined as:

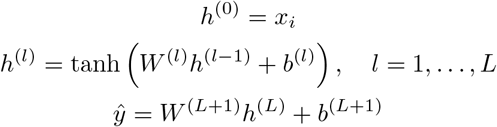

where *W* ^(*l*)^ and *b*^(*l*)^ denote the weights and biases of layer *l*. Depending on the experiment, *x*_*i*_ may correspond to the scGPT embedding, highly variable gene expression values, ion channel gene expression values, or one-hot cell type representation. The target vector *y*_*i*_ ∈ ℝ^8^ contains the eight percentile-normalized electrophysiological features. Our MLP architecture consisted of *L* = 2 tanh-activated hidden layers with 512 units each.

All models were implemented in PyTorch [33]. Models were trained using mini-batch gradient descent (SGD optimizer with a batch size of 25) for 1,500 epochs and tuned over the same set of hyperparameters (learning rate, weight decay, and dropout) using Optuna [34]. Mean squared error (MSE) was used as the loss function:

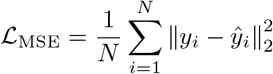

where *N* is the number of cells in the training set and ∥·∥_2_ is the *ℓ*^2^ norm of the vector.

For all experiments, we used stratified four-fold cross-validation (see S1 Fig). Cells of each type were evenly distributed across folds, and all models were trained and evaluated on identical splits, which were held constant across experiments. Hyperparameters were selected using nested three-fold cross-validation within each outer training fold, where the training data were further split into inner training and validation sets. The hyperparameter configuration that minimized the average validation loss at the final epoch across the three inner folds (out of the 100 configurations tested) was selected. Using these hyperparameters, a final model for a given outer fold was then trained on the full outer training set and evaluated on the held-out outer test set.

Model performance was quantified using the coefficient of determination (*R*^2^), computed separately for each electrophysiological feature:

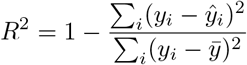

where 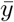 is the mean value of the target feature. Because all electrophysiological features were percentile-normalized across cells, 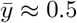 for every feature. Positive *R*^2^ values indicate predictive performance exceeding that of predicting this mean value, while negative values indicate worse-than-mean predictions.

While all experiments followed the general architecture and training and evaluation framework described above, individual tests incorporated task-specific modifications (e.g., the use of pretrained models or weight initialization), which are described in detail in the corresponding Results sections.

### AI use statement

ChatGPT and Perplexity were both used to proofread writing and assist in literature review. ChatGPT and Claude were used to boilerplate and proofread code. All AI-assisted text, references, and code were reviewed and verified by the authors, who take responsibility for the final content.

## Results

### Baseline comparison

As an initial comparison, we trained three MLPs, each with a distinct input representation: scGPT cell embeddings, the expression of the top 512 highly variable genes, or the expression of 328 ion channel-coding genes (refer Fig 2). All models were trained under the conditions described in the Materials and methods section above. Model performance, measured using *R*^2^, was compared against a cell type mean baseline, in which predictions were given by the mean electrophysiological features of each cell type as seen in the training data. Although all three models underperformed relative to the baseline, the model using scGPT embeddings achieved the highest performance among the three representations, matching or slightly exceeding baseline performance for a small subset of features.

**Fig 2.**
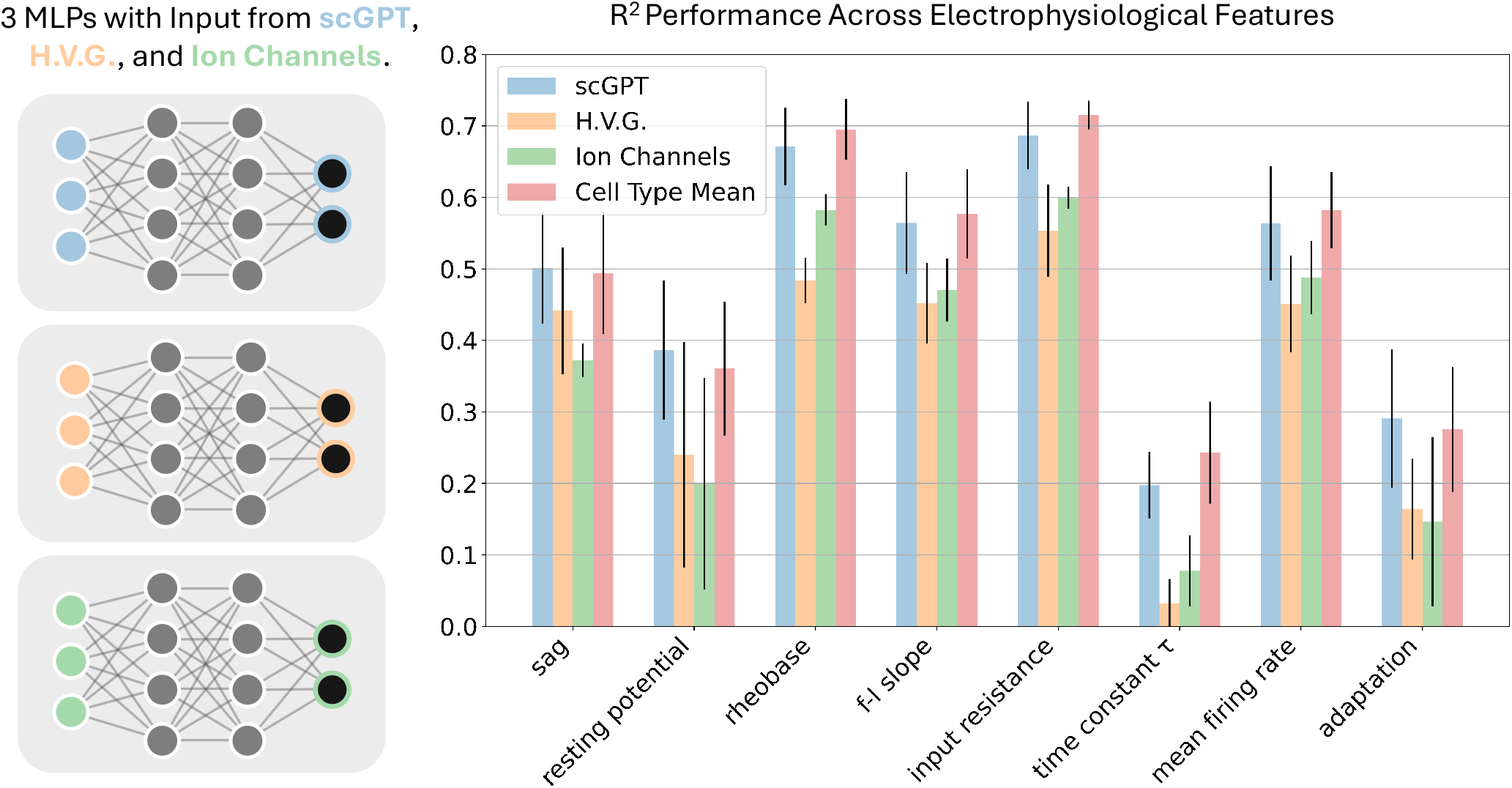
Model architectures and final epoch test R^2^ across electrophysiological features for the baseline comparison. A: Schematic of the three models, sharing the same architecture described in Materials and methods and varying only by size of the input layer. B: The mean prediction accuracies (*R*^2^) for every model across all percentile-transformed electrophysiological parameters. Shown in red is the result if a model were to predict cells in the test set based solely on cell type knowledge, using the mean feature values for each cell type from the training data. Error bars are standard deviations in the *R*^2^ values across the four folds of cross-validation.

### Comparison given cell type information

Given that the cell type means baseline outperformed all models in the initial test, we repeated the analysis while explicitly providing cell type information to each model. Specifically, we concatenate the transcriptomic representation *x*_*i*_ with a 15-dimensional one-hot encoding of cell type *c*_*i*_ in the input layer of each MLP:

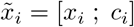

and use 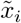 as input to the same network architecture, i.e. 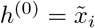.

Under this setting, the model using scGPT embeddings achieved the highest overall *R*^2^ across electrophysiological features and was the only model to outperform the cell type means baseline (Fig 3). In contrast, models based on highly variable genes or ion channel-coding genes continued to underperform, despite having access to cell type information.

**Fig 3.**
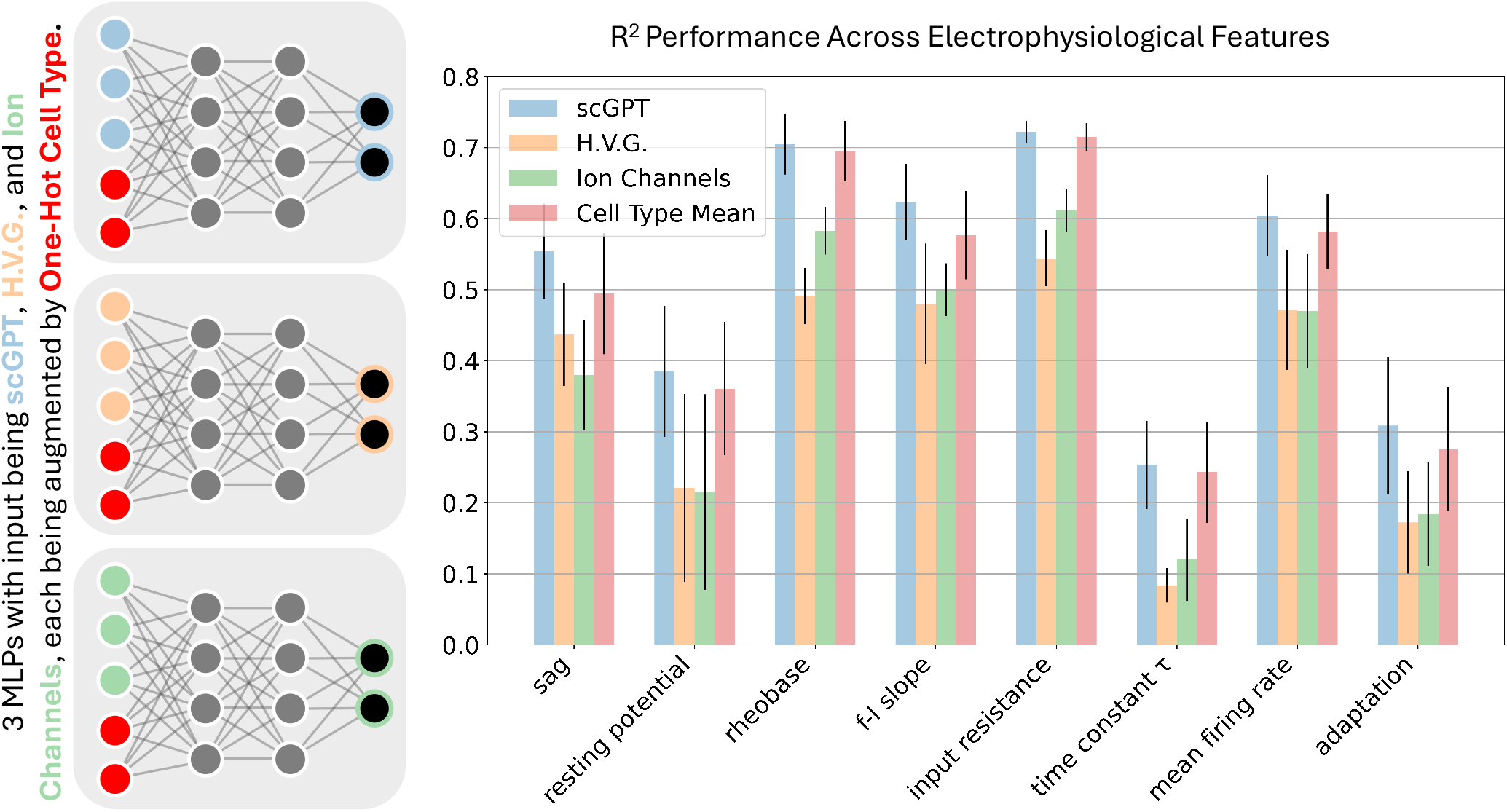
Model architectures and final epoch test R^2^ across electrophysiological features when given one-hot cell type input. A: Schematic of the three models, sharing the same architecture described in Materials and methods and varying only by size of the input layer. The input units in red represent the concatenated one-hot cell type. B: The mean prediction accuracies (*R*^2^) for every model across all percentile-transformed electrophysiological parameters. Shown in red is the result if a model were to predict cells in the test set based solely on cell type knowledge, using the mean feature values for each cell type from the training data. Error bars are standard deviations in the *R*^2^ values across the four folds of cross-validation.

### Comparison given cell type information and favorable initialization

While the models from the previous comparison all perform relatively similarly by the end of training, those trained on highly variable genes and ion channel genes fail to generalize to the test set, as seen in S10 Fig. To test if the performance of the networks given these inputs is consistently worse than the cell type mean, or merely a fault in our setup, we repeated the experiment, but initialize the sub-networks of the MLPs with weights from a “pretrained” network (sub-network being depicted using darker connections in Fig 4) that used only the one-hot cell type vector to predict electrophysiological features. That is, we first train a network using only cell type input:

**Fig 4.**
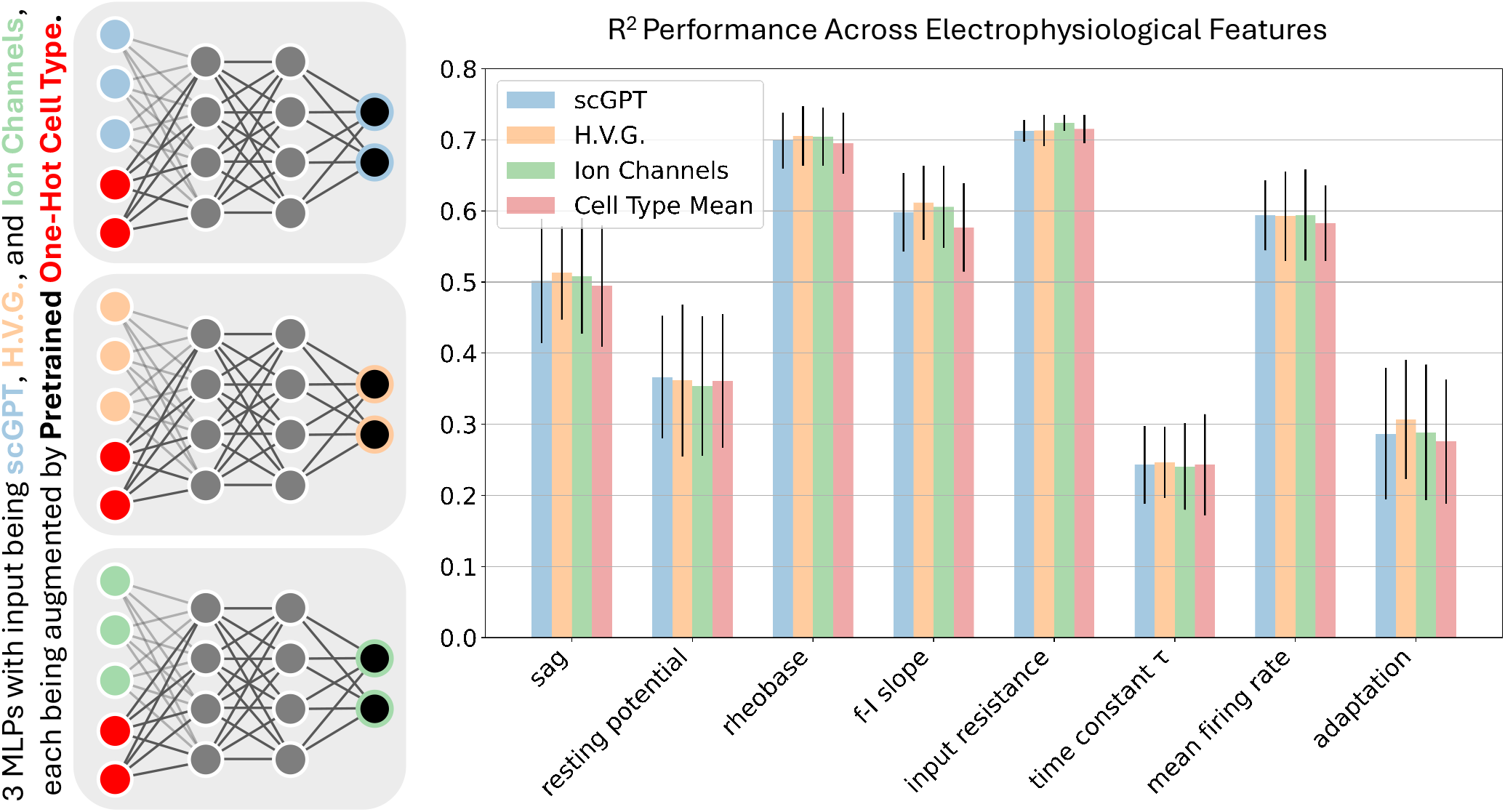
Model architectures and final epoch test R^2^ across electrophysiological features when granted one-hot cell type input and initialization. A: Schematic of the models, sharing the same architecture described in Materials and methods and varying only by size of the input layer. One-hot cell type input is shown in red. At the start of training, the weights of the larger model were set to match those of the one-hot only network, with the weights of the additional input features set to 0. B: The mean prediction accuracies (*R*^2^) for every model across all percentile-transformed electrophysiological parameters. Shown in red is the result if a model were to predict cells in the test set based solely on cell type knowledge, using the mean feature values for each cell type from the training data. Error bars are standard deviations in the *R*^2^ values across the four folds of cross-validation.

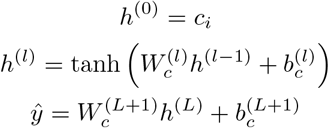

This pretrained network had 15 input units, *L* = 2 hidden layers of 512 units each, and eight output units (S11 Fig).

The learned weights and biases {*W*_*c*_, *b*_*c*_*}* are then used to initialize the corresponding parameters within the larger MLPs. Weights corresponding to additional input dimensions were initialized to zero. To avoid data leakage, we saved the trained networks from the inner folds of the best hyperparameter trial for the one-hot model. During the tuning of the larger network, initialization used the corresponding inner fold one-hot models.

For each outer fold, the final model was initialized using the one-hot network trained on the full outer training set (with hyperparameters selected during tuning) and then retrained on that same dataset as part of training the larger network, using the hyperparameters from the second (larger network) tuning procedure (S1 Fig).

Under this training regime, none of the models clearly outperform the others; all achieve performance at or very near the cell type mean baseline.

### Dual MLP fusion models

Our earlier results indicate that cell type identity provides a strong baseline for predicting electrophysiological features, while transcriptomic representations may carry additional, complementary information. However, directly concatenating cell type one-hot vectors with high-dimensional gene-based inputs can be difficult to optimize in small datasets and may obscure whether improvements arise from the transcriptomic signal or from model capacity and initialization. To more cleanly test whether transcriptomic representations add predictive information beyond cell type, we introduce a *dual MLP* model that combines two separately trained predictors.

The dual MLP architecture consists of two pretrained networks: one that takes experimental inputs,

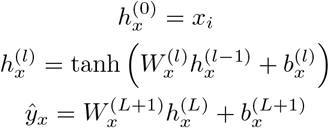

and another that takes one-hot cell type inputs,

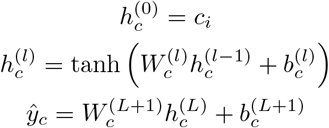

Their outputs are combined through a single linear output layer:

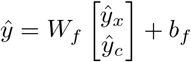

At the start of training, the dual MLP is initialized with the weights of the corresponding pretrained networks where applicable, and the final fusion layer is initialized to compute the average of the two subnetworks’ output features. The entire model is then trained using a new set of hyperparameters. By initializing in this way, we ensure that any performance gains arise from learned integration rather than favorable initialization.

In practice, we trained three networks of the same type used in the baseline comparison, along with the one-hot network, and then trained three dual MLPs. As in the earlier initialization comparison, we tuned the larger model by saving the trained networks from the inner folds of the best hyperparameter trial for both the one-hot and experimental models.

With this setup, the model using scGPT embeddings achieves its best performance, with it outperforming all others examined in our whole study (see Fig 5). In contrast, the models using highly variable genes and ion channel genes show little change compared to the previous test. Inspection of the test loss curves in S10 Fig reveals that the network using scGPT embeddings exhibits some learning, whereas the other two networks learn negligibly and fail to generalize.

**Fig 5.**
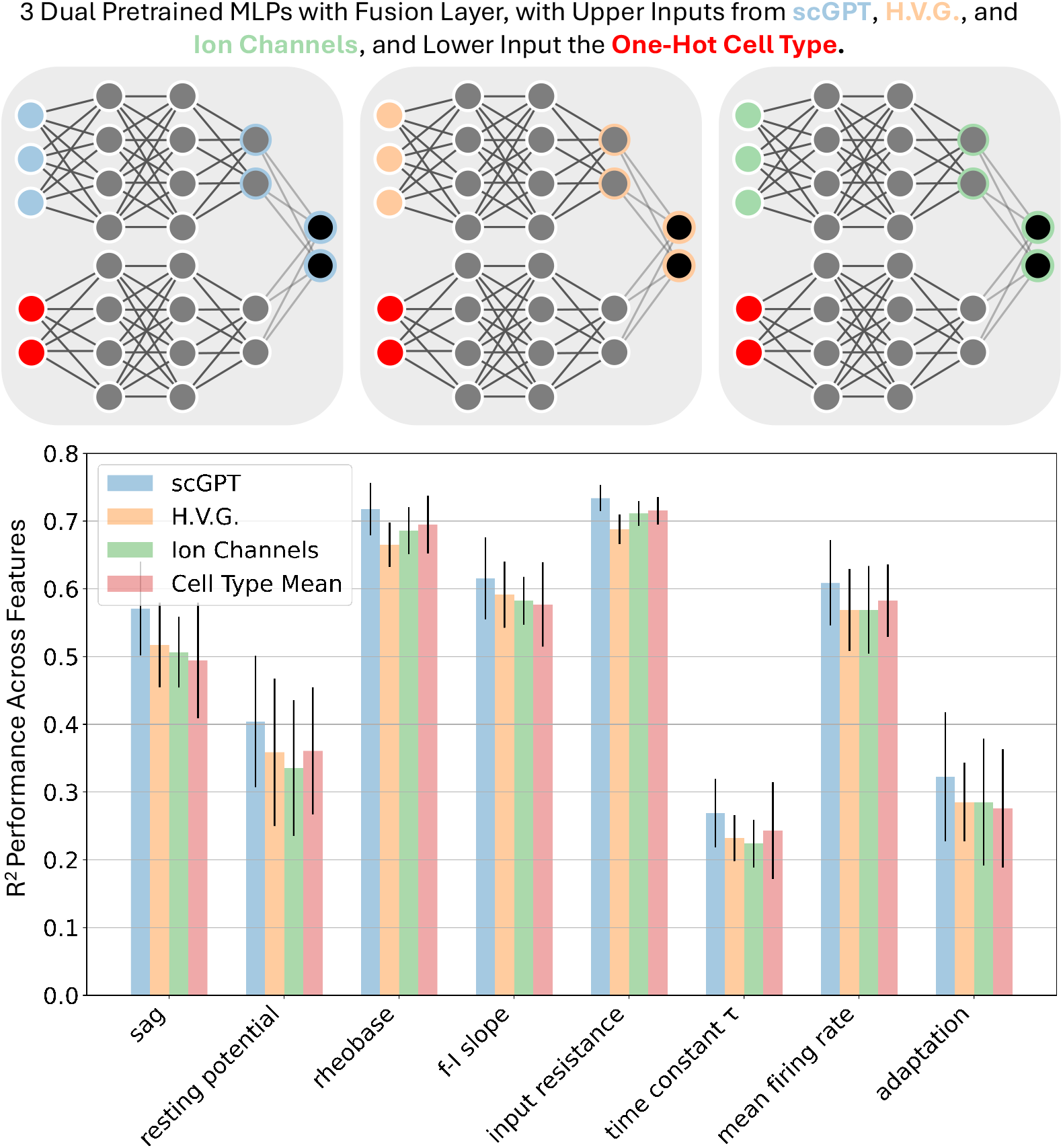
Model architectures and final epoch test R^2^ across electrophysiological features for the Dual MLP test. A: Schematic of the models, each consisting of two MLPs: one that takes experimental inputs and another that takes one-hot–encoded cell type inputs. The outputs of these MLPs are combined through a final linear layer. At the start of training, each MLP is initialized with weights from a model trained exclusively on its corresponding input type. The final linear layer is initialized to compute the mean of the two MLP outputs. The full combined network is then trained using the procedure outlined in the Materials and methods section. B: The mean prediction accuracies (*R*^2^) for every model across all percentile-transformed electrophysiological parameters. Shown in red is the result if a model were to predict cells in the test set based solely on cell type knowledge, using the mean feature values for each cell type from the training data. Error bars are standard deviations in the *R*^2^ values across the four folds of cross-validation.

## Discussion

Our initial baseline comparison suggests that single-cell transcriptomic data, in any continuous representation, is insufficient to outperform a simple cell type mean predictor. Highly variable genes, ion channel genes, and scGPT embeddings all contain biologically relevant information, but on their own they appear either too noisy, too weakly informative, or too difficult to exploit effectively in a dataset of only 495 cells. In contrast, transcriptomic cell type identity proved to be a remarkably strong summary statistic for neuronal physiology. Simply averaging electrophysiological properties within each cell type yielded performance that exceeded most transcriptome-based models.

At the same time, our results suggest that the scGPT representation remains useful when used to model variation within cell types rather than replace cell typing entirely. In our second experiment, naïvely concatenating the other transcriptomic representations with one-hot cell type labels did not consistently improve prediction. This indicates that simply providing both representations to a model is not sufficient for the model to discover how they should interact. Nevertheless, the scGPT embedding, when combined with cell type information, was able to exceed the performance of the baseline.

One interpretation of this result is that the scGPT embedding acts as a denoised and compressed representation of the full transcriptome. Rather than relying on sparse individual genes, scGPT likely captures broader co-expression programs and developmental gradients. This may explain why it consistently outperformed highly variable genes and ion channel genes. However, scGPT embeddings alone still did not surpass the cell type mean baseline, suggesting that current single-cell foundation model representations may be most valuable as complements to strong biological priors rather than standalone replacements.

Our initial observations are consistent with recent benchmarking studies of single-cell foundation models, which have shown that embeddings from models such as scGPT and Geneformer do not consistently outperform simpler baselines in low-data downstream tasks [35–37]. Across a range of evaluation settings, these studies find that relatively simple approaches such as linear modeling, Harmony, scVI, or even direct cell type labels often remain competitive with or superior to more complex pretrained representations. Taken together, these results suggest that the primary limitation may not be the representational quality of foundation models themselves, but rather the difficulty of extracting task-relevant signal from small datasets.

The relatively weak performance of ion channel genes alone was somewhat surprising given their mechanistic relationship to membrane potential dynamics. However, this result is consistent with prior literature suggesting that electrophysiological phenotypes cannot be explained solely by channel transcript abundance [4, 5, 38]. Intrinsic physiology is also shaped by dendritic morphology, neuromodulatory pathways, protein localization, and post-transcriptional regulation [39, 40]. Nuclear mRNA abundance is therefore only an indirect proxy for the functional density and localization of ion channels within a neuron.

Our experiments with initialized models further highlighted the information density of simple cell types. When transcriptomic representations were concatenated with cell type information and initialized from a model trained only on cell type, performance remained close to the original cell type baseline with negligible additional learning. The cell type representation provided such a strong inductive bias that the optimization procedure remained at a solution that ignored the additional transcriptomic information. This result also exposed a limitation of our modeling framework: although additional information was available, our training regime was not sufficient to encourage the models to make use of them.

To recover the information contained within the transcriptomic representations, we constructed Dual MLP models that processed cell type and transcriptomic inputs separately before combining them. These models yielded the strongest performance overall, with the best-performing model combining cell type with the scGPT embedding. This suggests that cell type and transcriptomic embeddings contain both redundant and complementary information, aligning with a newer view of transcriptomic identity as a combination of stable cell type programs and more transient state-dependent programs [41, 42]. Under this framework, one-hot cell types may capture the stable component of electrophysiological identity, while scGPT embeddings provide residual information about within-type variability. Consistent with this interpretation, we qualitatively observe that scGPT-based predictions most closely follow the distributions of the more populous cell types (S2–S9 Fig).

Several limitations should be considered when interpreting these findings. First, although 495 human Patch-seq neurons is substantial for a multimodal dataset, it remains very small relative to the dimensionality of transcriptomic data and the number of trainable parameters in deep neural networks. This imbalance likely limited the benefits of more complex models and contributed to the observed sensitivity of performance to architectural design and initialization, even when models were trained on identical inputs. Second, the modeled phenotype was itself incomplete: action potential features and latency were excluded in part because of their poor predictability in exploratory analyses, so performance on the retained features should not be interpreted as representative of the full electrophysiological phenotype. Third, aspects of the data generation and structure may have shaped our results. The weak performance of models using highly variable genes and ion channel gene sets may reflect limitations of the snRNA-seq modality, which may not fully capture within-cell-type transcriptional variability. In particular, snRNA-seq can miss aspects of alternative splicing, spatially localized transcription, and temporal delays between transcriptional state and phenotype. Additionally, cells were nested within donors varying in pathology and demographics, and cross-validation folds were stratified by cell type rather than donor, leaving open the possibility that donor-level effects contributed to the reported performance. Finally, our analysis was restricted to layer 1 interneurons and layer 2/3 glutamatergic neurons in human cortex, a composition spanning both broad neuronal classes and finer transcriptomic types. Aggregate performance may therefore reflect separation between broad classes as much as variation among finer types, and their respective contributions were not separately resolved here. It also remains unclear whether the same conclusions would generalize to deeper cortical layers, other brain regions, or larger mouse Patch-seq datasets.

Overall, our findings suggest that foundation model embeddings do contain meaningful information about electrophysiology, but that this information is most useful when layered on top of biologically interpretable cell type structure. In small multimodal neuroscience datasets, simple biological abstractions remain surprisingly competitive. Rather than replacing existing transcriptomic taxonomies, pretrained transcriptomic representations appear most effective when used to model continuous variation around these discrete categories.

Future work could extend these findings in several directions. Foremost, moving beyond electrophysiological prediction to neuronal morphology would help determine whether transcriptomic and foundation model embeddings capture structural as well as functional properties of neurons. Secondly, applying these approaches to larger mouse Patch-seq datasets would enable more expressive modeling and better resolution of within-cell-type variation, addressing the primary limitation of small sample size in the present study. Third, benchmarking alternative foundation model embeddings such as UCE, both alone and in combination with structured biological priors, would help clarify the effects of different pretraining strategies in encoding complementary or redundant information [43]. Lastly, our results suggest a more general question about how best to combine discrete cell type annotations with continuous transcriptomic embeddings. Further exploration could involve more explicit interaction mechanisms such as gating or mixture-of-experts models.

## Data and code availability

All code used for this study is available at https://github.com/harsh1ls/TransformerEphysPrediction. Data and model files are additionally available from https://doi.org/10.5281/zenodo.20500624.

## Acknowledgments

We wish to thank the Allen Institute founder, Paul G. Allen, for his vision, encouragement, and support. S.M. has been in part supported by NSF 2424124, 2223725, NIH R01EB029813, and RF1DA055669 grants.

## References

1. Lipovsek M, Bardy C, Cadwell CR, Hadley K, Kobak D, Tripathy SJ. Patch-seq: Past, Present, and Future. Journal of Neuroscience. 2021;41(5):937–46. Available from: https://www.jneurosci.org/content/41/5/937. arXiv: https://www.jneurosci.org/content/41/5/937.full.pdf. doi:10.1523/JNEUROSCI.1653-20.2020.

2. Fuzik J, Zeisel A, Máté Z, Calvigioni D, Yanagawa Y, Szabó G, et al. Integration of electrophysiological recordings with single-cell RNA-seq data identifies neuronal subtypes. Nature Biotechnology. 2016 Feb;34(2):175–83. Available from: https://doi.org/10.1038/nbt.3443. doi:10.1038/nbt.3443.

3. Földy C, Darmanis S, Aoto J, Malenka RC, Quake SR, Südhof TC. Single-cell RNAseq reveals cell adhesion molecule profiles in electrophysiologically defined neurons. Proceedings of the National Academy of Sciences. 2016;113(35):E5222–31. Available from: https://www.pnas.org/doi/abs/10.1073/pnas.1610155113. arXiv: https://www.pnas.org/doi/pdf/10.1073/pnas.1610155113. doi:10.1073/pnas.1610155113.

4. Tripathy SJ, Toker L, Li B, Crichlow CL, Tebaykin D, Mancarci BO, et al. Transcriptomic correlates of neuron electrophysiological diversity. PLOS Computational Biology. 2017 Oct;13(10):1–28. Available from: https://doi.org/10.1371/journal.pcbi.1005814. doi:10.1371/journal.pcbi.1005814.

5. Bomkamp C, Tripathy SJ, Bengtsson Gonzales C, Hjerling-Leffler J, Craig AM, Pavlidis P. Transcriptomic correlates of electrophysiological and morphological diversity within and across excitatory and inhibitory neuron classes. PLOS Computational Biology. 2019 Jun;15(6):1–33. Available from: https://doi.org/10.1371/journal.pcbi.1007113. doi:10.1371/journal.pcbi.1007113.

6. Nandi A, Chartrand T, Van Geit W, Buchin A, Yao Z, Lee SY, et al. Single-neuron models linking electrophysiology, morphology, and transcriptomics across cortical cell types. Cell Reports. 2022 Aug;40(6). Available from: https://doi.org/10.1016/j.celrep.2022.111176. doi:10.1016/j.celrep.2022.111176.

7. Cadwell CR, Palasantza A, Jiang X, Berens P, Deng Q, Yilmaz M, et al. Electrophysiological, transcriptomic and morphologic profiling of single neurons using Patch-seq. Nature Biotechnology. 2016 Feb;34(2):199–203. Available from: https://doi.org/10.1038/nbt.3445. doi:10.1038/nbt.3445.

8. Tasic B, Menon V, Nguyen TN, Kim TK, Jarsky T, Yao Z, et al. Adult mouse cortical cell taxonomy revealed by single cell transcriptomics. Nature Neuroscience. 2016 Feb;19(2):335–46. Available from: https://doi.org/10.1038/nn.4216. doi:10.1038/nn.4216.

9. Muñoz-Manchado AB, Bengtsson Gonzales C, Zeisel A, Munguba H, Bekkouche B, Skene NG, et al. Diversity of Interneurons in the Dorsal Striatum Revealed by Single-Cell RNA Sequencing and PatchSeq. Cell Reports. 2018 Aug;24(8):2179–90.e7. Available from: https://doi.org/10.1016/j.celrep.2018.07.053. doi:10.1016/j.celrep.2018.07.053.

10. Tasic B, Yao Z, Graybuck LT, Smith KA, Nguyen TN, Bertagnolli D, et al. Shared and distinct transcriptomic cell types across neocortical areas. Nature. 2018 Nov;563(7729):72–8. Available from: https://doi.org/10.1038/s41586-018-0654-5. doi:10.1038/s41586-018-0654-5.

11. Gouwens NW, Sorensen SA, Baftizadeh F, Budzillo A, Lee BR, Jarsky T, et al. Integrated Morphoelectric and Transcriptomic Classification of Cortical GABAergic Cells. Cell. 2020;183(4):935–53.e19. Available from: https://www.sciencedirect.com/science/article/pii/S009286742031254X. doi:10.1016/j.cell.2020.09.057.

12. Scala F, Kobak D, Bernabucci M, Bernaerts Y, Cadwell CR, Castro JR, et al. Phenotypic variation of transcriptomic cell types in mouse motor cortex. Nature. 2021 Oct;598(7879):144–50. Available from: https://doi.org/10.1038/s41586-020-2907-3. doi:10.1038/s41586-020-2907-3.

13. Yao Z, Velthoven CTJv, Nguyen TN, Goldy J, Sedeno-Cortes AE, Baftizadeh F, et al. A taxonomy of transcriptomic cell types across the isocortex and hippocampal formation. Cell. 2021;184(12):3222–41.e26. Available from: https://www.sciencedirect.com/science/article/pii/S0092867421005018. doi:10.1016/j.cell.2021.04.021.

14. Gala R, Budzillo A, Baftizadeh F, Miller J, Gouwens N, Arkhipov A, et al. Consistent cross-modal identification of cortical neurons with coupled autoencoders. Nature Computational Science. 2021 Feb;1(2):120–7. Available from: https://doi.org/10.1038/s43588-021-00030-1. doi:10.1038/s43588-021-00030-1.

15. Furumichi K, Kojima Y, Nomura S, Shimamura T. A deep generative model integrating single-cell time-frequency characteristics transformed from electrophysiological data with transcriptomic features. bioRxiv. 2024 Jan:2024.03.29.587341. Available from: http://biorxiv.org/content/early/2024/04/01/2024.03.29.587341.abstract. doi:10.1101/2024.03.29.587341.

16. Schneider A, Azabou M, McDougall-Vigier L, Parks DF, Ensley S, Bhaskaran-Nair K, et al. Transcriptomic cell type structures in vivo neuronal activity across multiple timescales. Cell Reports. 2023;42(4):112318. Available from: https://www.sciencedirect.com/science/article/pii/S2211124723003297. doi:10.1016/j.celrep.2023.112318.

17. Hodge RD, Bakken TE, Miller JA, Smith KA, Barkan ER, Graybuck LT, et al. Conserved cell types with divergent features in human versus mouse cortex. Nature. 2019 Sep;573(7772):61–8. Available from: https://doi.org/10.1038/s41586-019-1506-7. doi:10.1038/s41586-019-1506-7.

18. Chartrand T, Dalley R, Close J, Goriounova NA, Lee BR, Mann R, et al. Morphoelectric and transcriptomic divergence of the layer 1 interneuron repertoire in human versus mouse neocortex. Science. 2023;382(6667):eadf0805. Available from: https://www.science.org/doi/abs/10.1126/science.adf0805. arXiv: https://www.science.org/doi/pdf/10.1126/science.adf0805. doi:10.1126/science.adf0805.

19. Mathys H, Davila-Velderrain J, Peng Z, Gao F, Mohammadi S, Young JZ, et al. Single-cell transcriptomic analysis of Alzheimer’s disease. Nature. 2019 Jun;570(7761):332–7. Available from: https://doi.org/10.1038/s41586-019-1195-2. doi:10.1038/s41586-019-1195-2.

20. Velmeshev D, Schirmer L, Jung D, Haeussler M, Perez Y, Mayer S, et al. Single-cell genomics identifies cell type–specific molecular changes in autism. Science. 2019;364(6441):685–9. Available from: https://www.science.org/doi/abs/10.1126/science.aav8130. arXiv: https://www.science.org/doi/pdf/10.1126/science.aav8130. doi:10.1126/science.aav8130.

21. Bardy C, van den Hurk M, Kakaradov B, Erwin JA, Jaeger BN, Hernandez RV, et al. Predicting the functional states of human iPSC-derived neurons with single-cell RNA-seq and electrophysiology. Molecular Psychiatry. 2016 Nov;21(11):1573–88. Available from: https://doi.org/10.1038/mp.2016.158. doi:10.1038/mp.2016.158.

22. Szalata A, Hrovatin K, Becker S, Tejada-Lapuerta A, Cui H, Wang B, et al. Transformers in single-cell omics: a review and new perspectives. Nature Methods. 2024 Aug;21(8):1430–43. Available from: https://doi.org/10.1038/s41592-024-02353-z. doi:10.1038/s41592-024-02353-z.

23. Baek S, Song K, Lee I. Single-cell foundation models: bringing artificial intelligence into cell biology. Experimental & Molecular Medicine. 2025 Oct;57(10):2169–81. Available from: https://doi.org/10.1038/s12276-025-01547-5. doi:10.1038/s12276-025-01547-5.

24. Cui H, Wang C, Maan H, Pang K, Luo F, Duan N, et al. scGPT: toward building a foundation model for single-cell multi-omics using generative AI. Nature Methods. 2024 Aug;21(8):1470–80. Available from: https://doi.org/10.1038/s41592-024-02201-0. doi:10.1038/s41592-024-02201-0.

25. Berg J, Sorensen SA, Ting JT, Miller JA, Chartrand T, Buchin A, et al. Human neocortical expansion involves glutamatergic neuron diversification. Nature. 2021 Oct;598(7879):151–8. Available from: https://doi.org/10.1038/s41586-021-03813-8. doi:10.1038/s41586-021-03813-8.

26. Lake BB, Codeluppi S, Yung YC, Gao D, Chun J, Kharchenko PV, et al. A comparative strategy for single-nucleus and single-cell transcriptomes confirms accuracy in predicted cell-type expression from nuclear RNA. Scientific Reports. 2017 Jul;7(1):6031. Available from: https://doi.org/10.1038/s41598-017-04426-w. doi:10.1038/s41598-017-04426-w.

27. Bakken TE, Hodge RD, Miller JA, Yao Z, Nguyen TN, Aevermann B, et al. Single-nucleus and single-cell transcriptomes compared in matched cortical cell types. PLOS ONE. 2018 Dec;13(12):1–24. Available from: https://doi.org/10.1371/journal.pone.0209648. doi:10.1371/journal.pone.0209648.

28. Slyper M, Porter CBM, Ashenberg O, Waldman J, Drokhlyansky E, Wakiro I, et al. A single-cell and single-nucleus RNA-Seq toolbox for fresh and frozen human tumors. Nature Medicine. 2020 May;26(5):792–802. Available from: https://doi.org/10.1038/s41591-020-0844-1. doi:10.1038/s41591-020-0844-1.

29. Stuart T, Butler A, Hoffman P, Hafemeister C, Papalexi E, III WMM, et al. Comprehensive Integration of Single-Cell Data. Cell. 2019;177:1888–902. Available from: https://doi.org/10.1016/j.cell.2019.05.031. doi:10.1016/j.cell.2019.05.031.

30. Allen Institute for Brain Science. Allen Cell Types Database Technical Whitepaper: Electrophysiology; 2017. Available from: https://celltypes.brain-map.org/.

31. Wolf FA, Angerer P, Theis FJ. SCANPY: large-scale single-cell gene expression data analysis. Genome Biology. 2018 Feb;19(1):15. Available from: https://doi.org/10.1186/s13059-017-1382-0. doi:10.1186/s13059-017-1382-0.

32. Seal RL, Braschi B, Gray K, McClay J, Tweedie S, Bruford EA. Genenames.org: the HGNC and PGNC resources in 2026. Nucleic Acids Research. 2025 Nov;54(D1):D1098-107. eprint: https://academic.oup.com/nar/article-pdf/54/D1/D1098/65494932/gkaf1229.pdf. Available from: https://doi.org/10.1093/nar/gkaf1229. doi:10.1093/nar/gkaf1229.

33. Paszke A, Gross S, Massa F, Lerer A, Bradbury J, Chanan G, et al. PyTorch: An Imperative Style, High-Performance Deep Learning Library; 2019. Available from: https://arxiv.org/abs/1912.01703. arXiv:1912.01703.

34. Akiba T, Sano S, Yanase T, Ohta T, Koyama M. Optuna: A Next-generation Hyperparameter Optimization Framework. In: Proceedings of the 25th ACM SIGKDD International Conference on Knowledge Discovery & Data Mining. KDD ‘19. New York, NY, USA: Association for Computing Machinery; 2019. p. 2623–2631. Available from: https://doi.org/10.1145/3292500.3330701. doi:10.1145/3292500.3330701.

35. Kedzierska KZ, Crawford L, Amini AP, Lu AX. Zero-shot evaluation reveals limitations of single-cell foundation models. Genome Biology. 2025 Apr;26(1):101. Available from: https://doi.org/10.1186/s13059-025-03574-x. doi:10.1186/s13059-025-03574-x.

36. Wong DR, Hill AS, Moccia R. Simple controls exceed best deep learning algorithms and reveal foundation model effectiveness for predicting genetic perturbations. Bioinformatics. 2025 Jun;41(6):btaf317. eprint: https://academic.oup.com/bioinformatics/article-pdf/41/6/btaf317/63308149/btaf317.pdf. Available from: https://doi.org/10.1093/bioinformatics/btaf317. doi:10.1093/bioinformatics/btaf317.

37. Ahlmann-Eltze C, Huber W, Anders S. Deep-learning-based gene perturbation effect prediction does not yet outperform simple linear baselines. Nature Methods. 2025 Aug;22(8):1657–61. Available from: https://doi.org/10.1038/s41592-025-02772-6. doi:10.1038/s41592-025-02772-6.

38. Goaillard JM, Marder E. Ion Channel Degeneracy, Variability, and Covariation in Neuron and Circuit Resilience. Annual Review of Neuroscience. 2021;44(Volume 44, 2021):335–57. Type: Journal Article. Available from: https://www.annualreviews.org/content/journals/10.1146/annurev-neuro-092920-121538. doi:10.1146/annurev-neuro-092920-121538.

39. Schulz DJ, Temporal S, Barry DM, Garcia ML. Mechanisms of voltage-gated ion channel regulation: from gene expression to localization. Cellular and Molecular Life Sciences. 2008 Jul;65(14):2215–31. Available from: https://doi.org/10.1007/s00018-008-8060-z. doi:10.1007/s00018-008-8060-z.

40. Cembrowski M, Bachman J, Wang L, Sugino K, Shields B, Spruston N. Spatial Gene-Expression Gradients Underlie Prominent Heterogeneity of CA1 Pyramidal Neurons. Neuron. 2016 Jan;89(2):351–68. Available from: https://doi.org/10.1016/j.neuron.2015.12.013. doi:10.1016/j.neuron.2015.12.013.

41. Trapnell C. Defining cell types and states with single-cell genomics. Genome Research. 2015;25(10):1491–8. eprint: http://genome.cshlp.org/content/genome/25/10/1491.full.pdf. Available from: http://genome.cshlp.org/content/genome/25/10/1491. doi:10.1101/gr.190595.115.

42. Olmstead JA, King LE, Bloodgood BL. Intersection of transient cell states with stable cell types in hippocampus. 2025 Dec. Available from: http://dx.doi.org/10.7554/eLife.109663.1. doi:10.7554/elife.109663.1.

43. Rosen Y, Roohani Y, Agarwal A, Samotorčan L, Tabula Sapiens Consortium, Quake SR, et al. Universal Cell Embeddings: A Foundation Model for Cell Biology. bioRxiv. 2023 Jan:2023.11.28.568918. Available from: http://biorxiv.org/content/early/2023/11/29/2023.11.28.568918.abstract. doi:10.1101/2023.11.28.568918.

