## Supplementary Figures for "The Unreasonable Effectiveness of Cell Types in Describing Neuronal Physiological Features"

### Supporting information

##### S1 Fig.

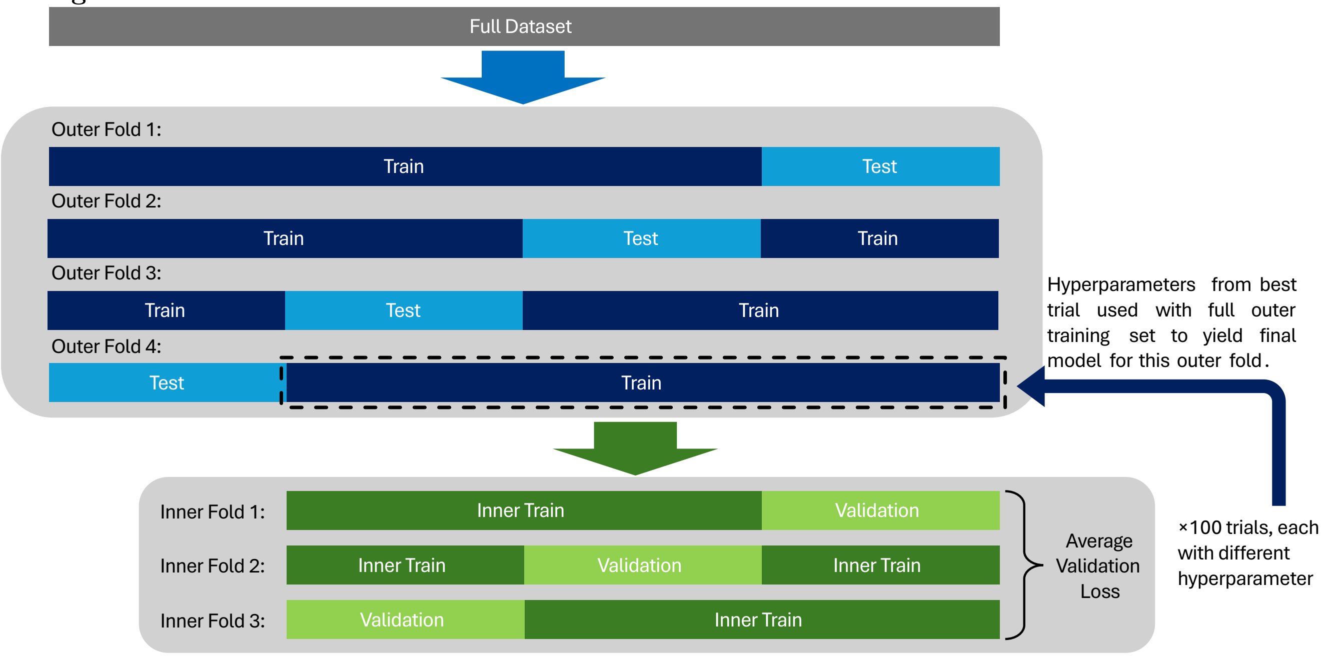


**Visualization of the nested cross-validation loop.** This study used stratified four-fold nested cross-validation. The full dataset comprising of 495 samples is split into four unique folds, ensuring that the cell types in both the training and test data were representative of the overall dataset. Prior to training models on each training set, hyperparameter tuning was performed. To tune the hyperparameters, the training data of each outer fold was further partitioned into three stratified sub-folds, creating inner training and validation sets. During each tuning trial, a set of hyperparameters was selected, used to train a model on the inner training set, and then evaluated on the corresponding validation set for each of the three inner folds. The validation performance across the three inner folds was averaged to compute an overall score (average loss) for the hyperparameter set. This process was repeated for 100 tuning trials, and the hyperparameter set with the lowest average loss on the validation datasets was chosen. Once the optimal hyperparameters were determined, the model was trained using these parameters on the full outer training set and evaluated on the unseen test data.

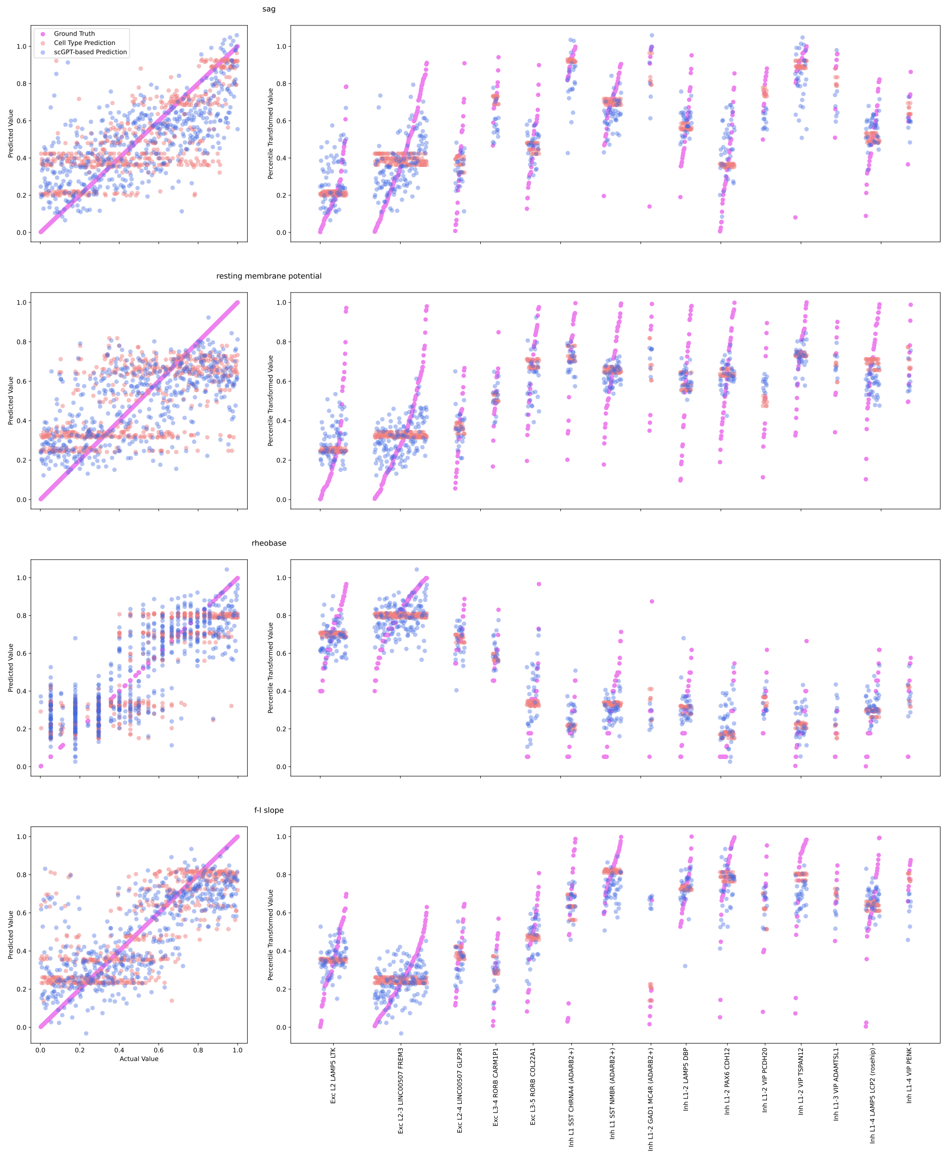


**S2 Fig. Actual versus predicted for first four electrophysiological features in the baseline comparison.** Right: Distributions across cells grouped by cell type, along with the predictions from the MLP using the scGPT embeddings and the predictions from cell type mean. Results from test data across all folds of cross-validation are shown (i.e. every cell in the dataset is plotted). Left: Same as right but without distinguishing cell type.

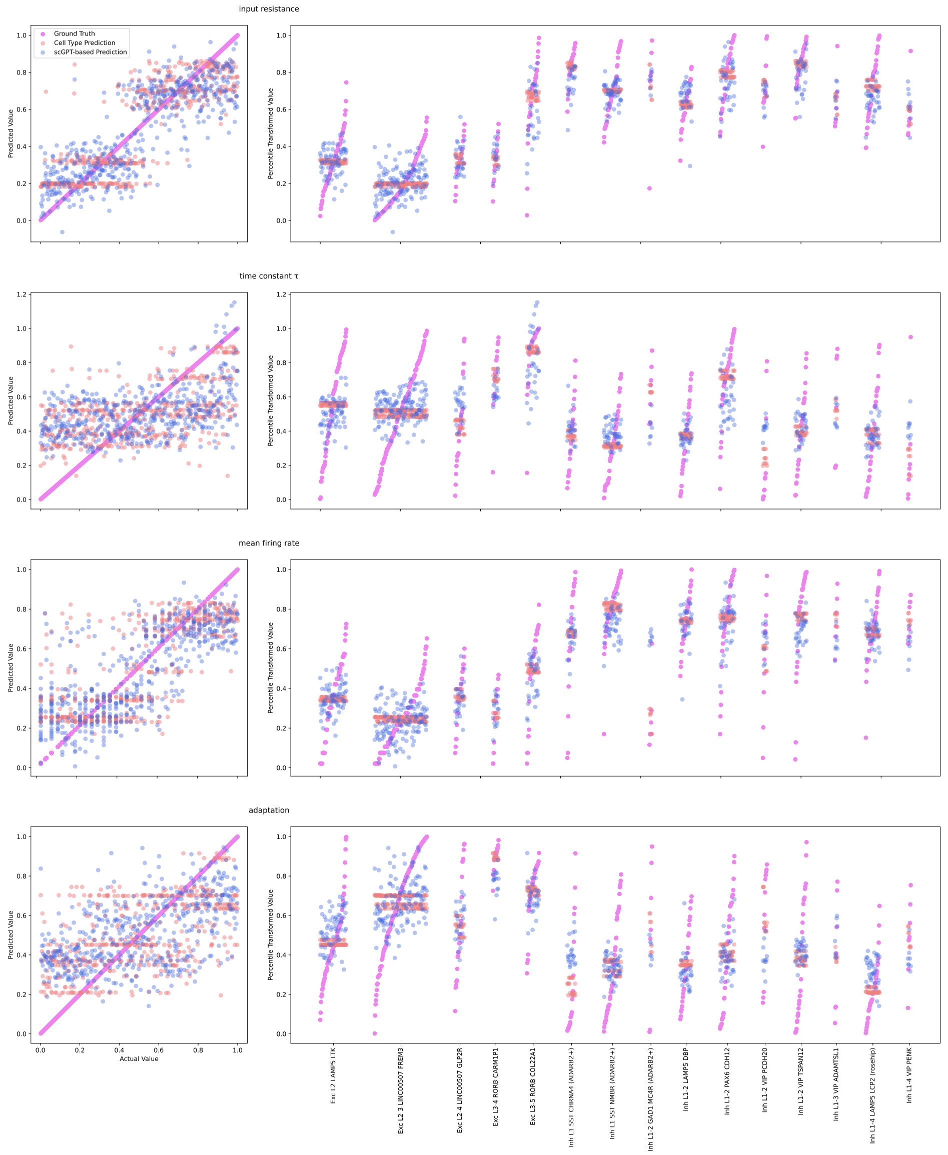


**S3 Fig. Actual versus predicted for last four electrophysiological features in the baseline comparison.** Right: Distributions of predictions across cells grouped by cell type, along with the predictions from the MLP using the scGPT embeddings and the predictions from cell type mean. Results from test data across all folds of cross-validation are shown (i.e. every cell in the dataset is plotted). Left: Same as right but without distinguishing cell type.

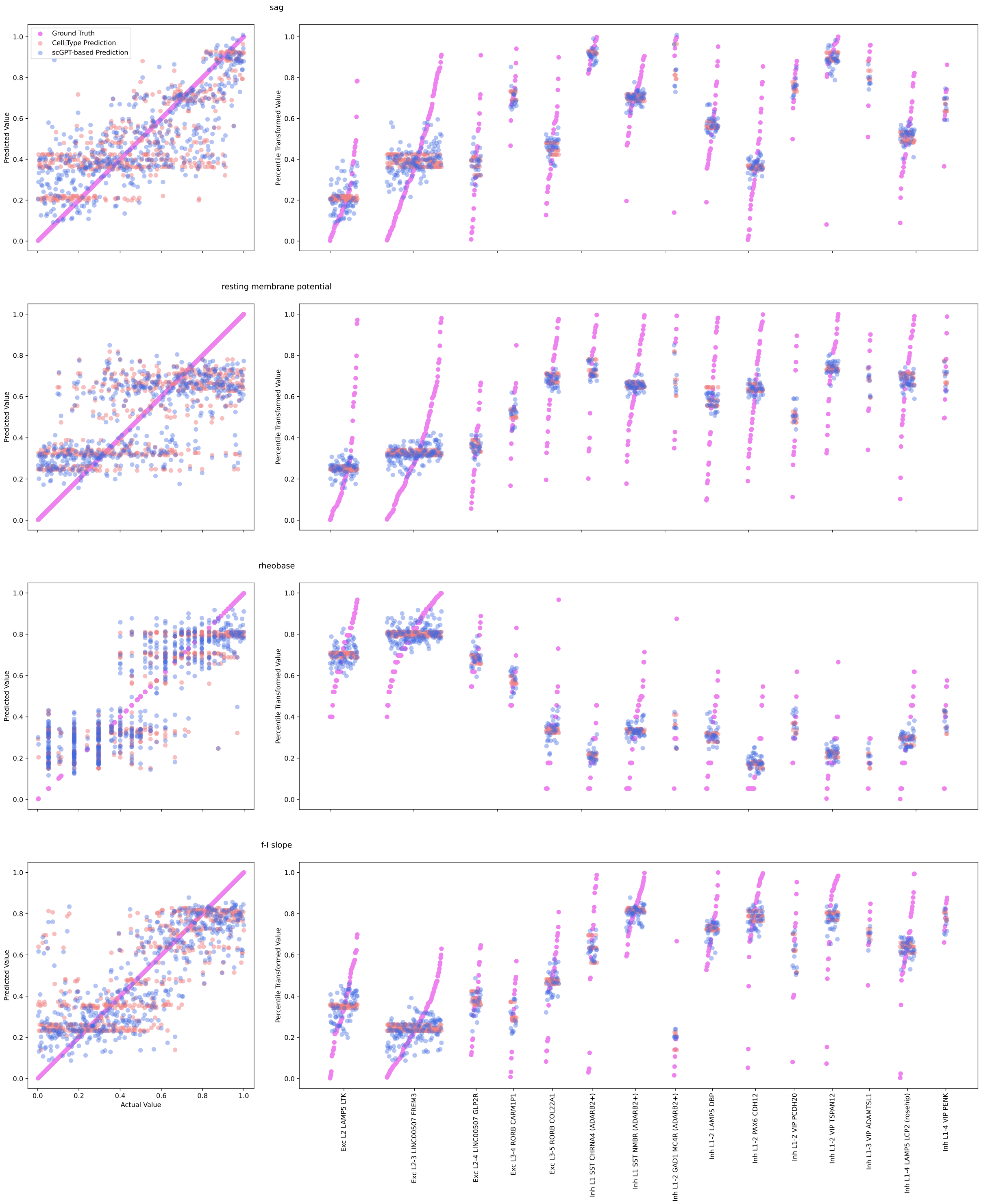


**S4 Fig.**
**Actual versus predicted for first four electrophysiological features in the comparison given cell type information.**

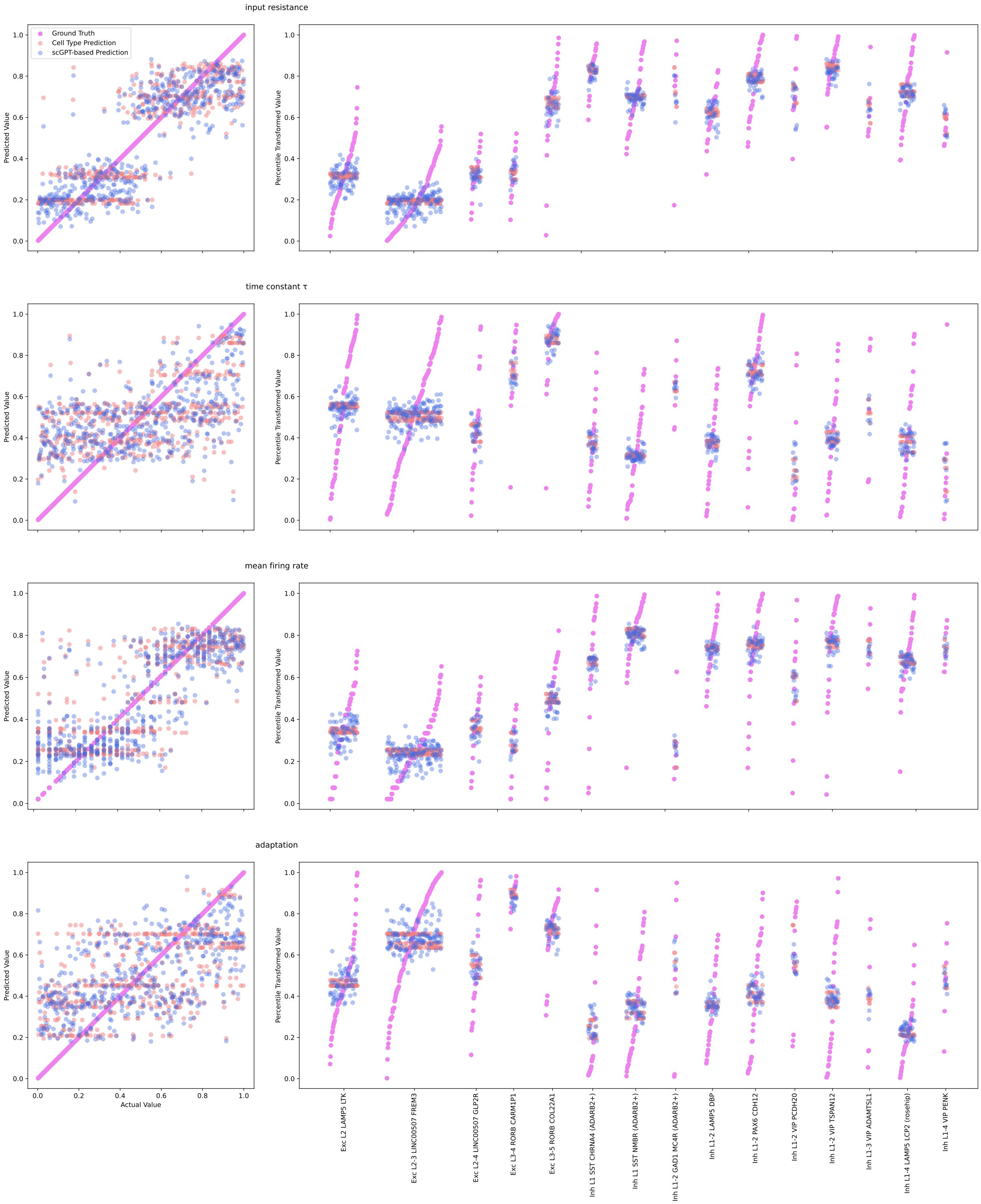


**S5 Fig.**
**Actual versus predicted for last four electrophysiological features in the comparison given cell type information.**

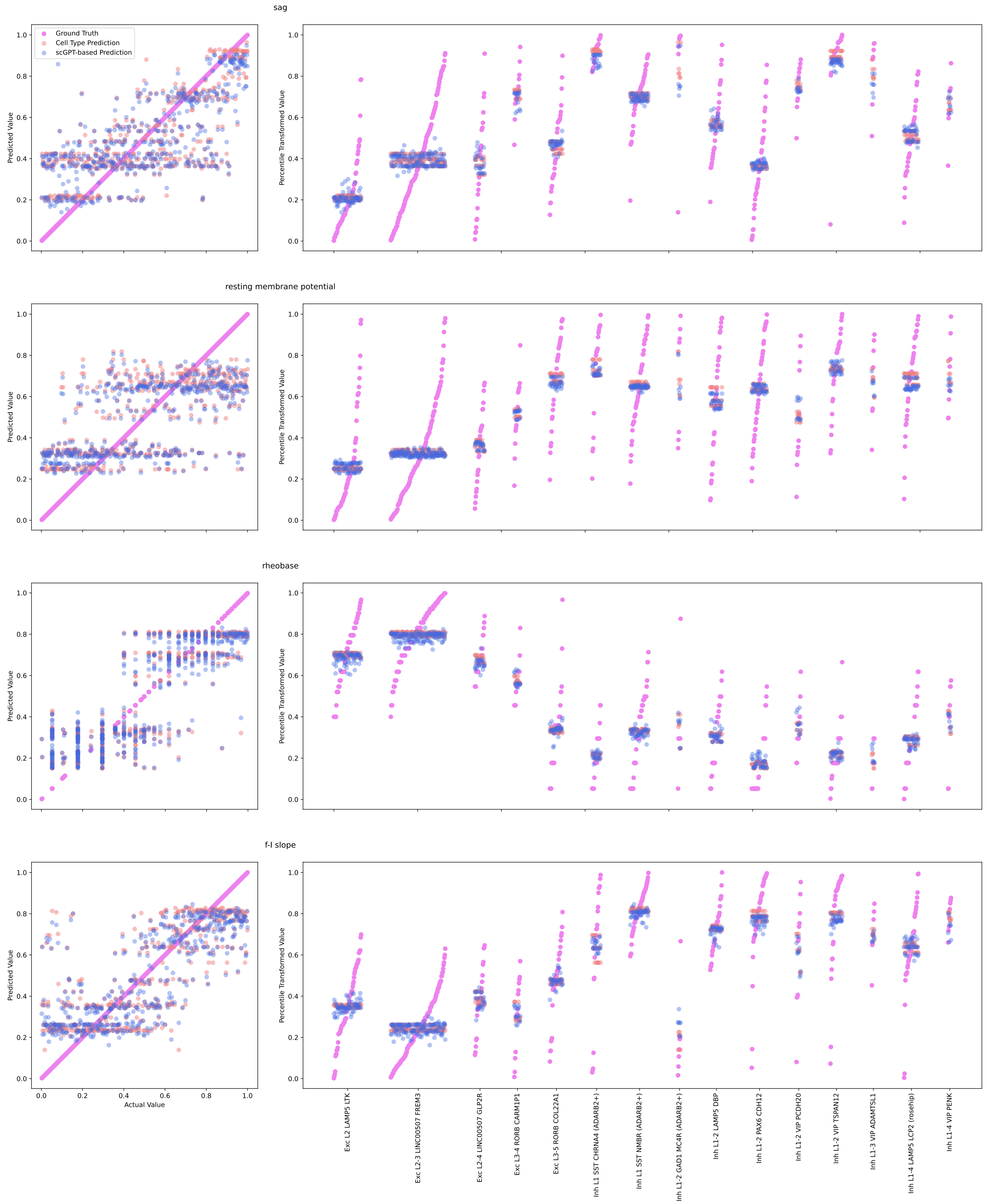


**S6 Fig.**

**Actual versus predicted for first four electrophysiological features in the comparison given cell type information and favorable initialization.**

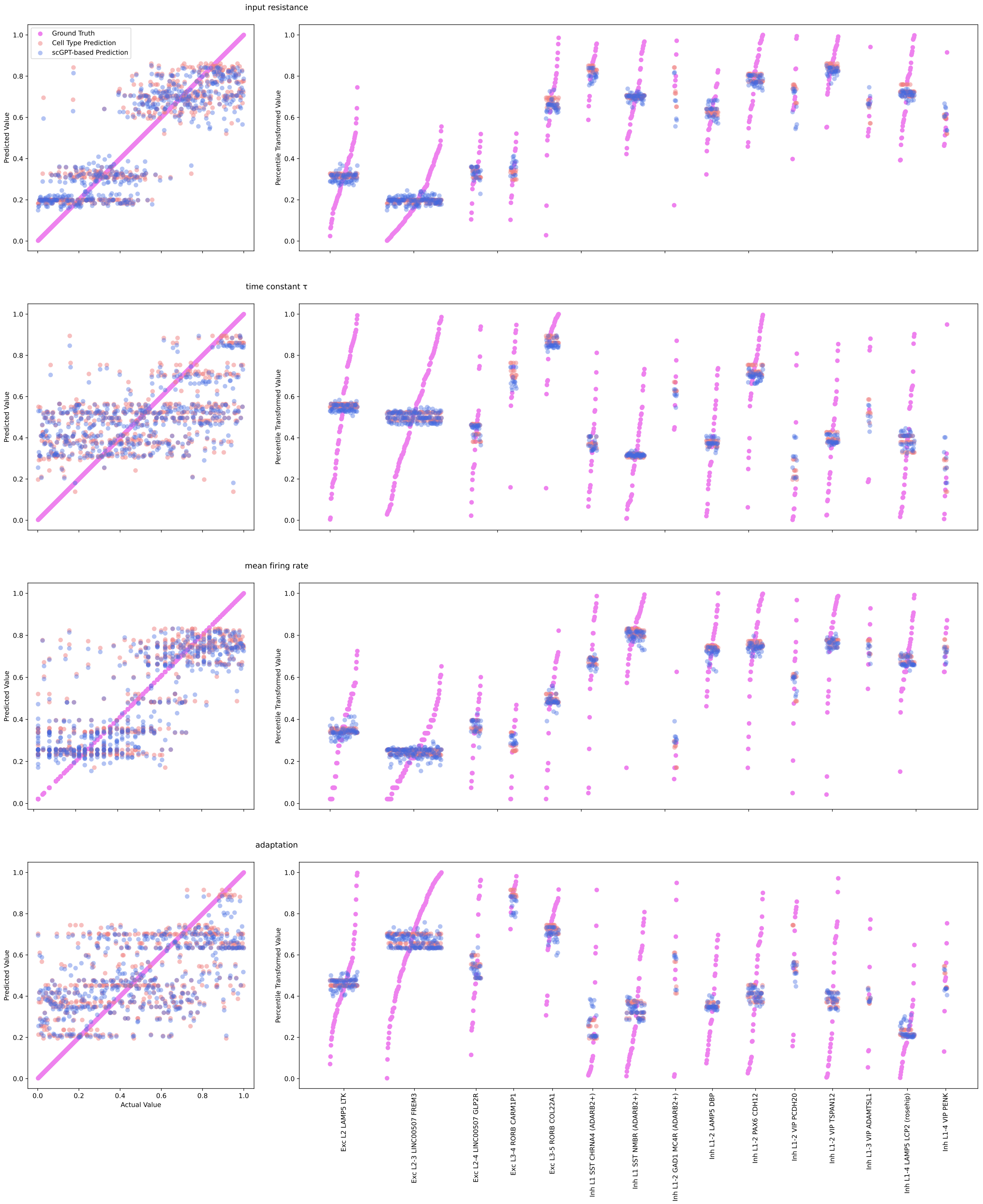


**S7 Fig.**

**Actual versus predicted for last four electrophysiological features in the comparison given cell type information and favorable initialization.**

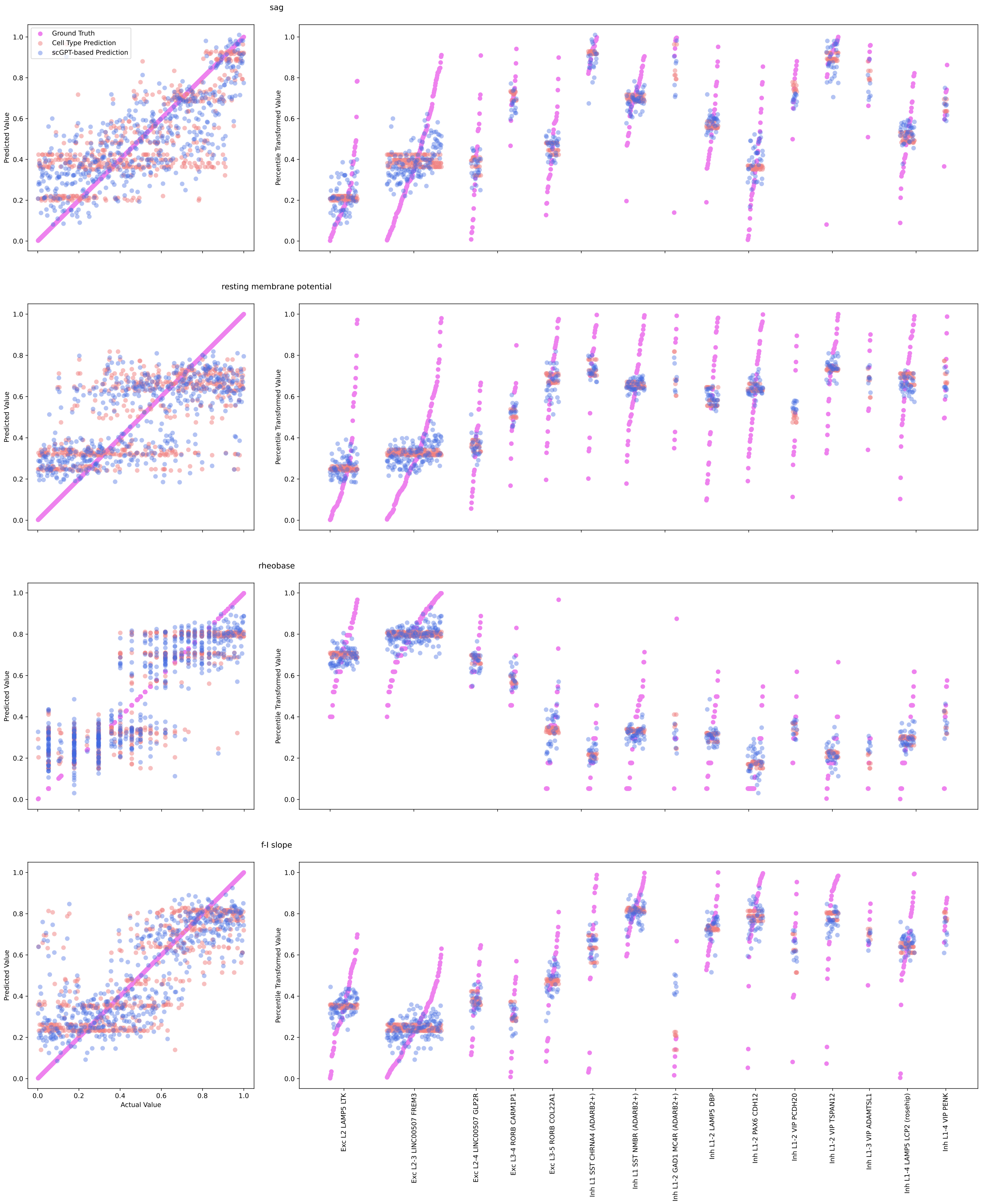


**S8 Fig.**
**Actual versus predicted for first four electrophysiological features in the Dual MLP comparison.**

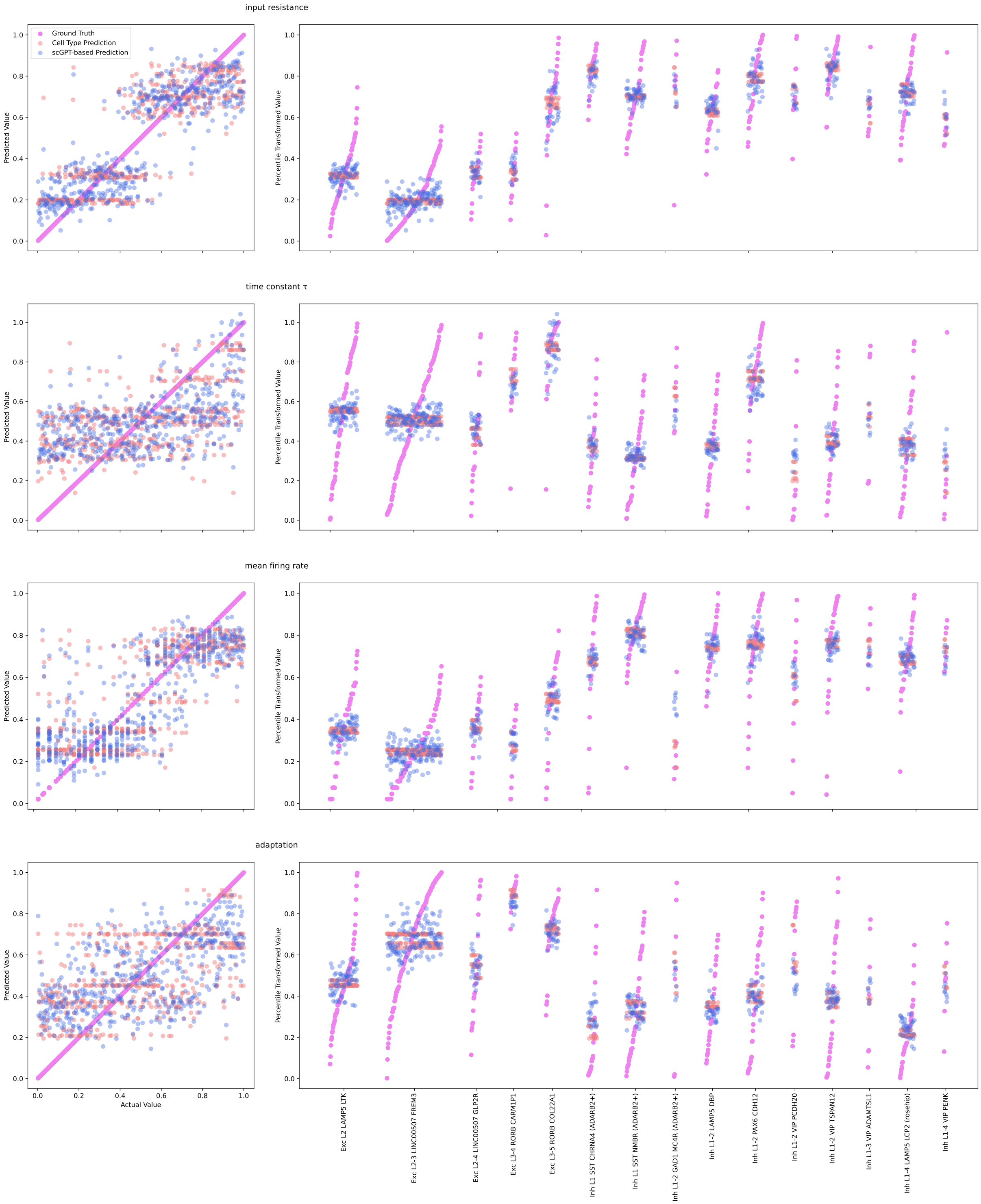


**S9 Fig.**
**Actual versus predicted for last four electrophysiological features in the Dual MLP comparison.**

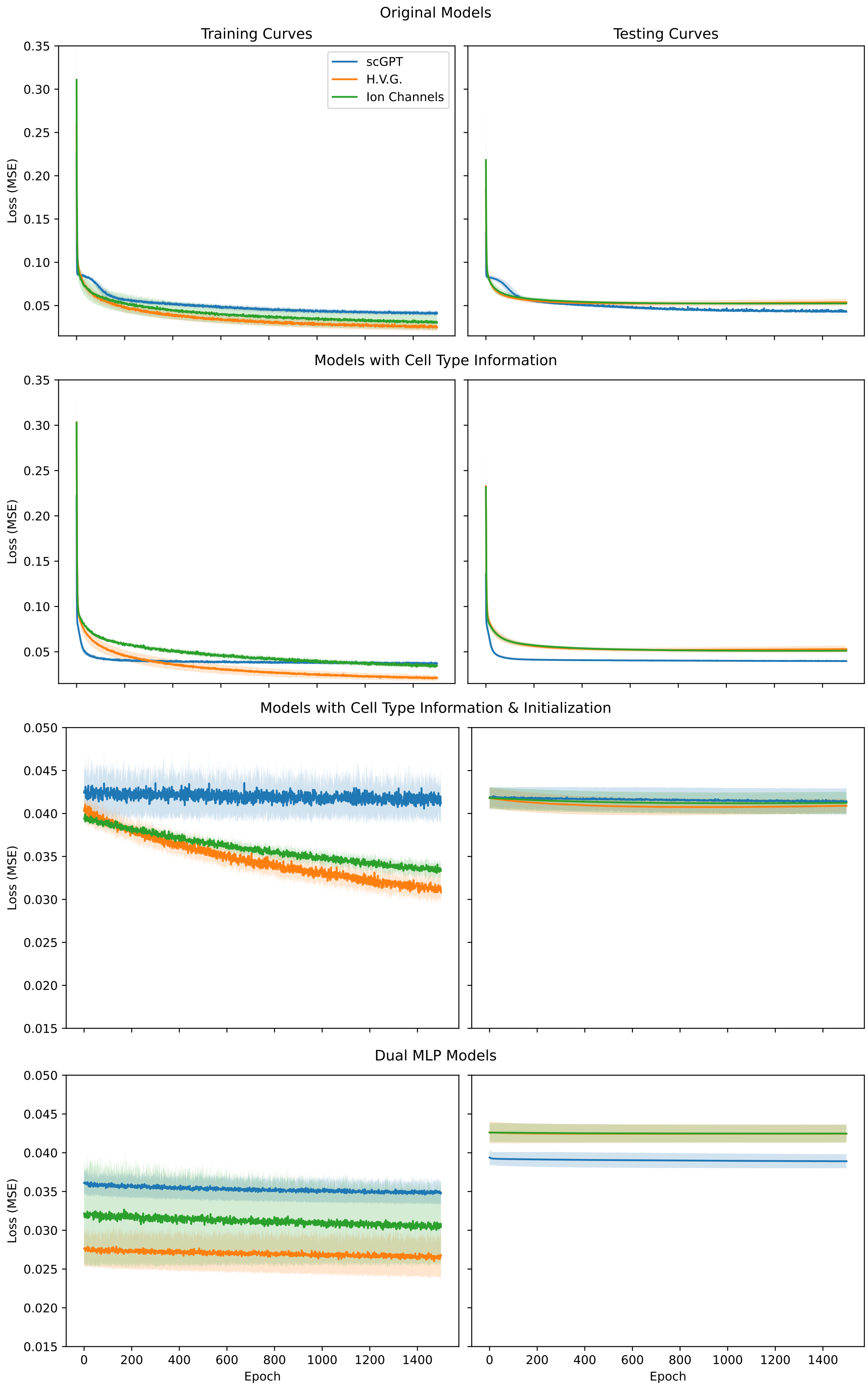


**S10 Fig. Learning curves for all MLPs.** Error bars are standard deviations across the four folds of cross-validation.

**S11 Fig.**

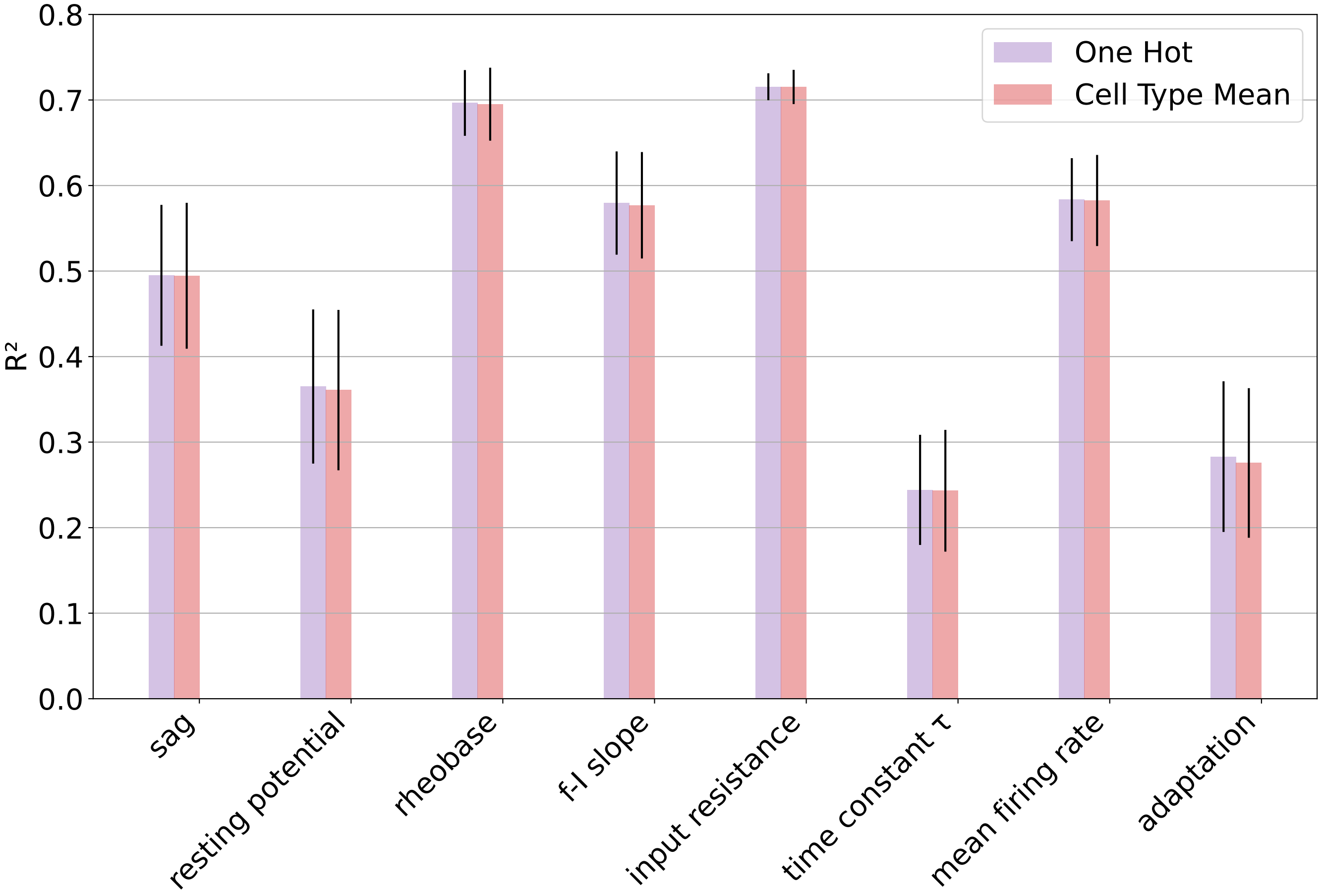


**Performance of MLP taking in exclusively one-hot cell type against that of cell type mean predictions.** The mean prediction accuracies ($R^2$) for the MLP trained only using one-hot cell type across all percentile-transformed electrophysiological parameters. Shown in red is the result if predictions simply used the mean feature values for each cell type from the training data. Error bars are standard deviations in the $R^2$ values across the four folds of cross-validation.

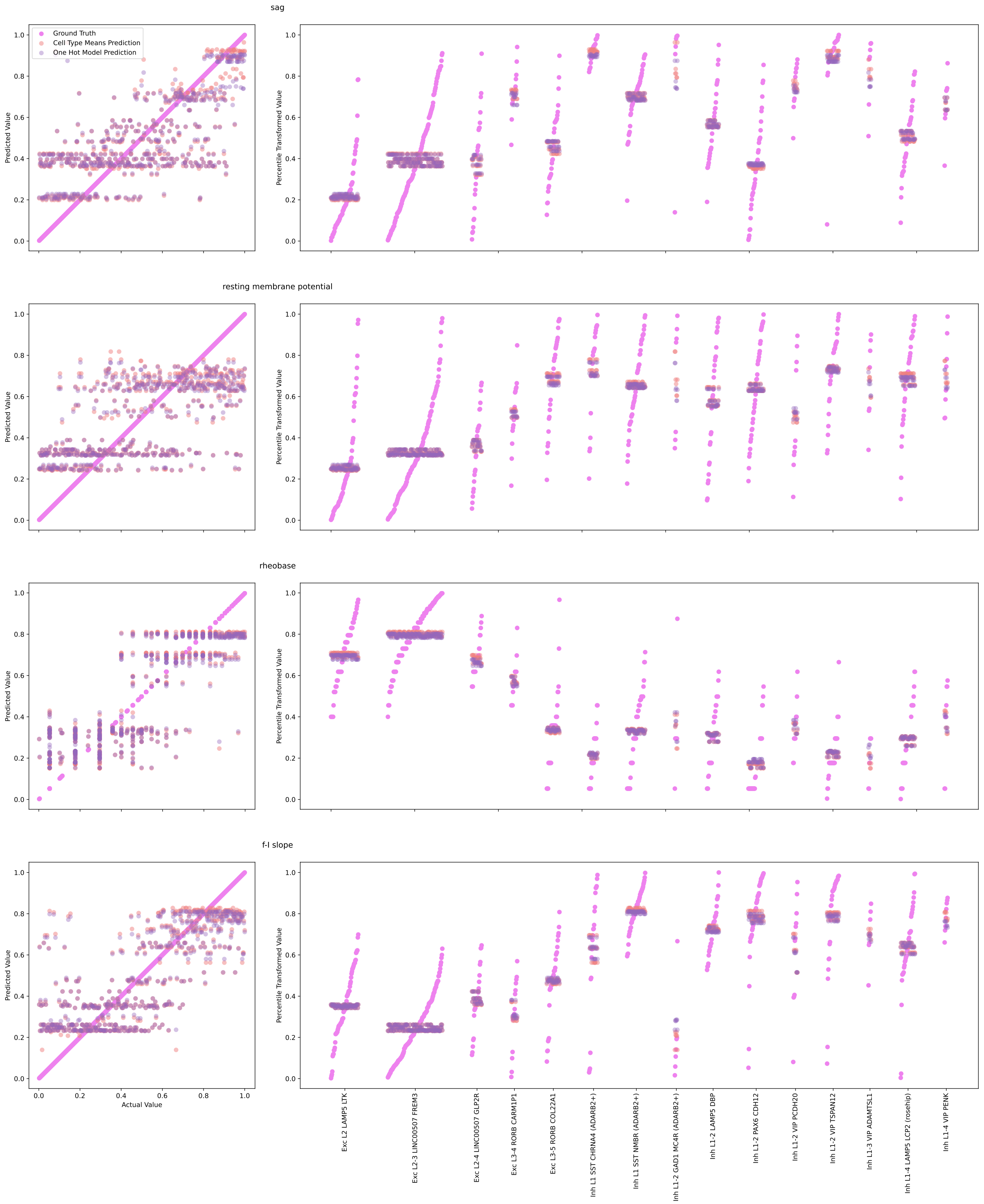


**S12 Fig.**
**Predictions from the pretrained one-hot MLP compared to those of the cell type means for the first four electrophysiological features.**

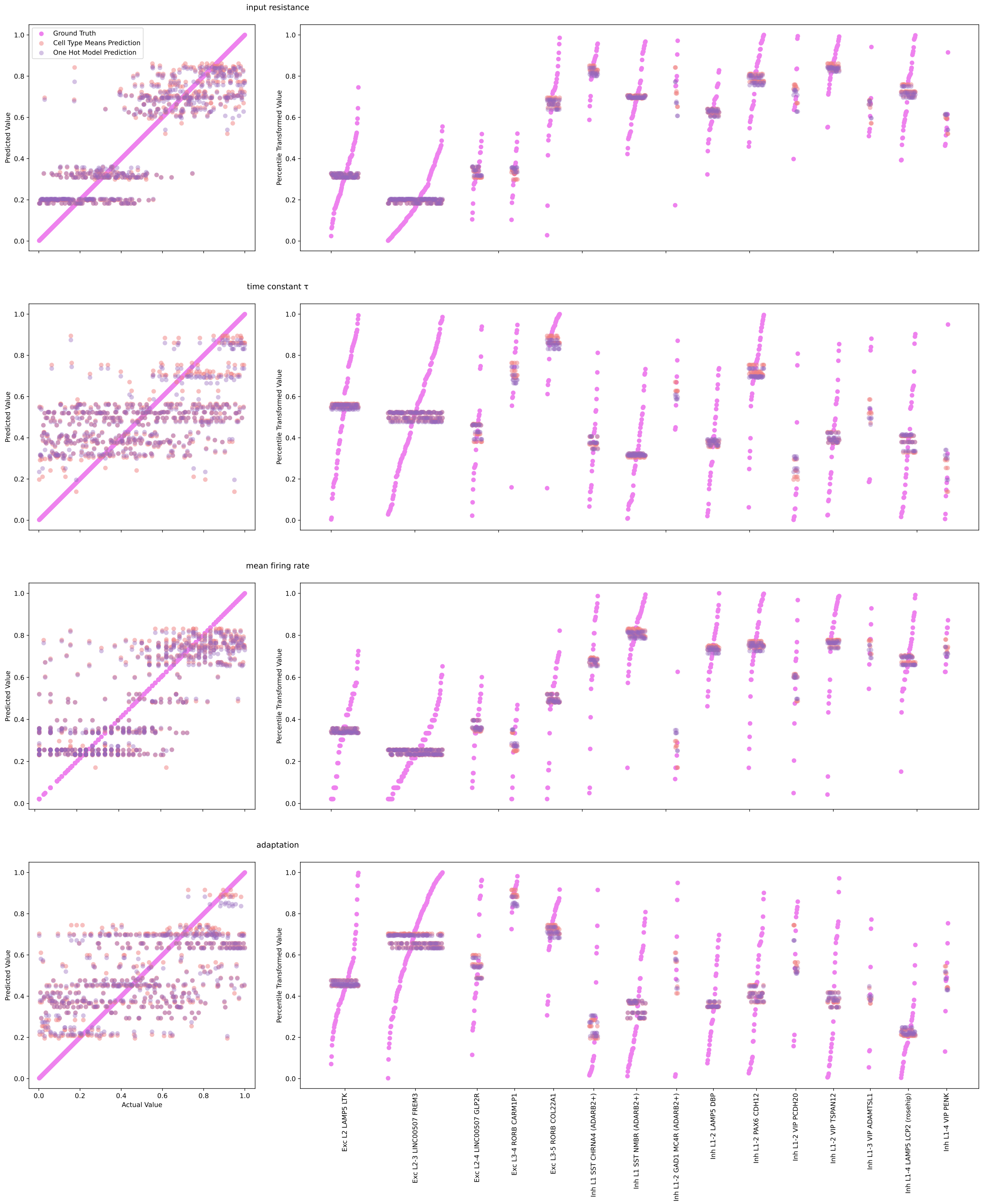


**S13 Fig.**
**Predictions from the pretrained one-hot MLP compared to those of the cell type means for the last four electrophysiological features.**

**S14 Fig.**

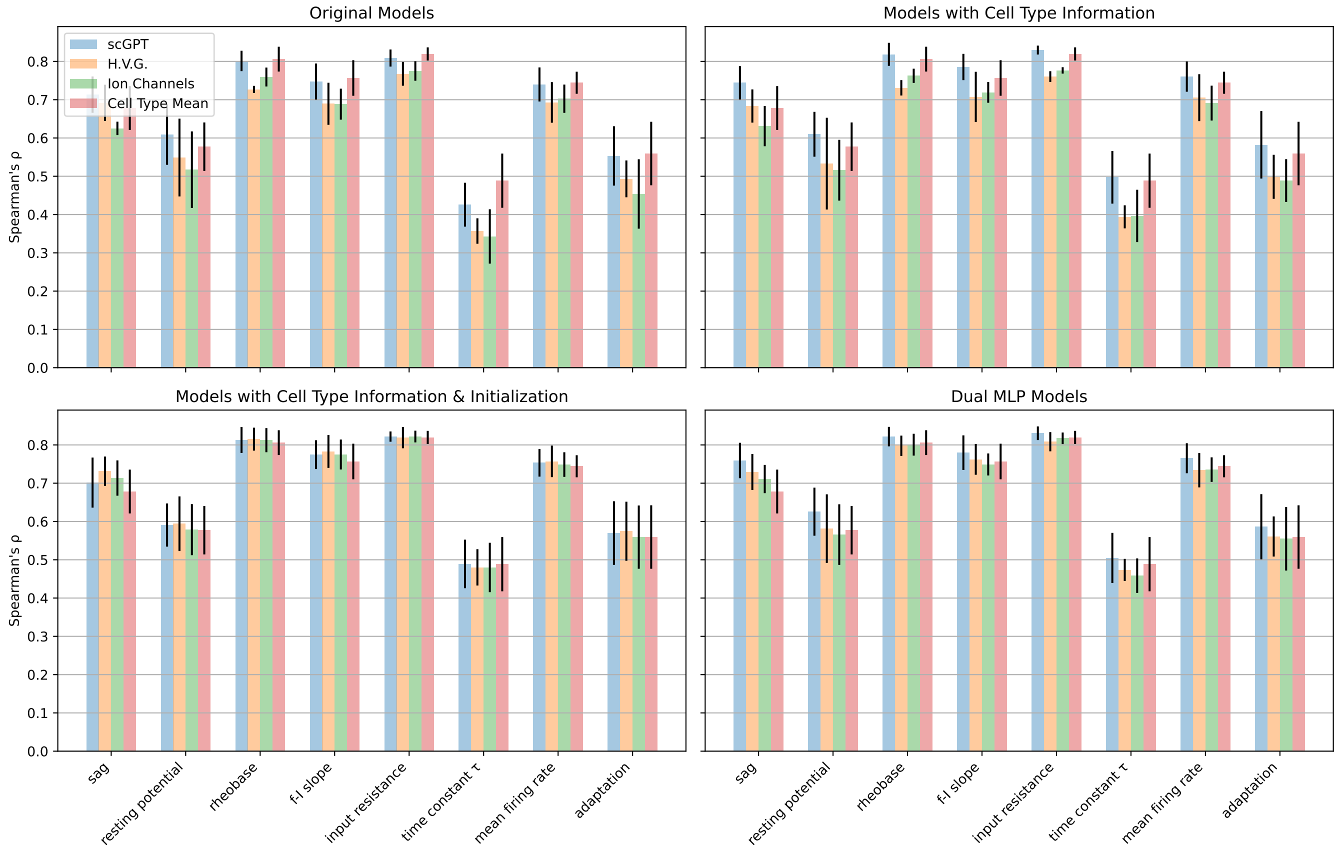


**Performance of models across all tests and features measured using Spearman's rank correlation coefficient.** Spearman's $\rho$ for every model across all percentile-transformed electrophysiological parameters. Shown in red is the result if a model were to predict cells in the test set based solely on cell type knowledge, using the mean feature values for each cell type from the training data. Error bars are standard deviations in the $\rho$ values across the four folds of cross-validation.

**S15 Fig.**

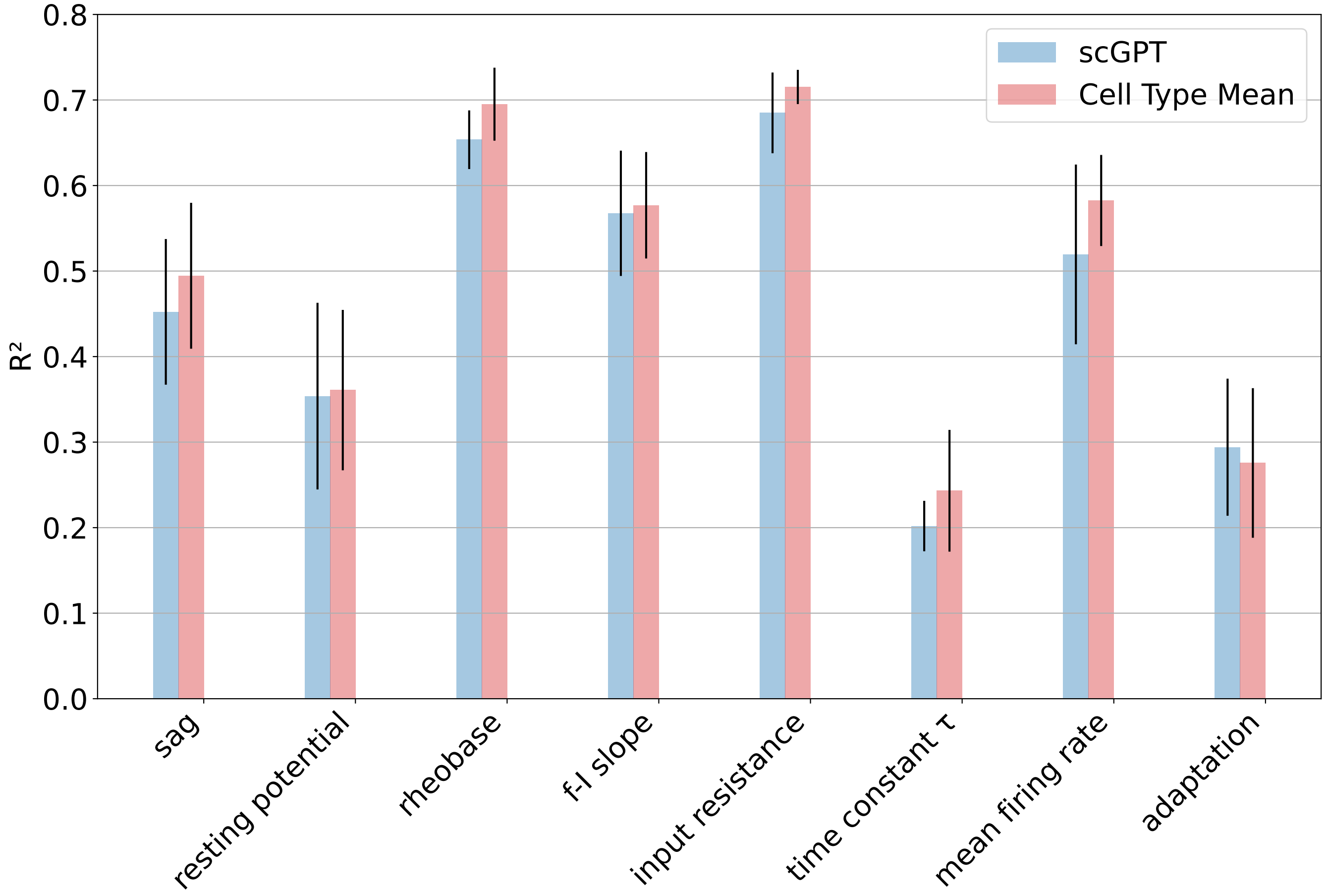


**Performance of MLPs given solely scGPT embeddings as input, but predicting only one output feature at a time.** Error bars are standard deviations across the four folds of cross-validation.

**S16 Fig.**

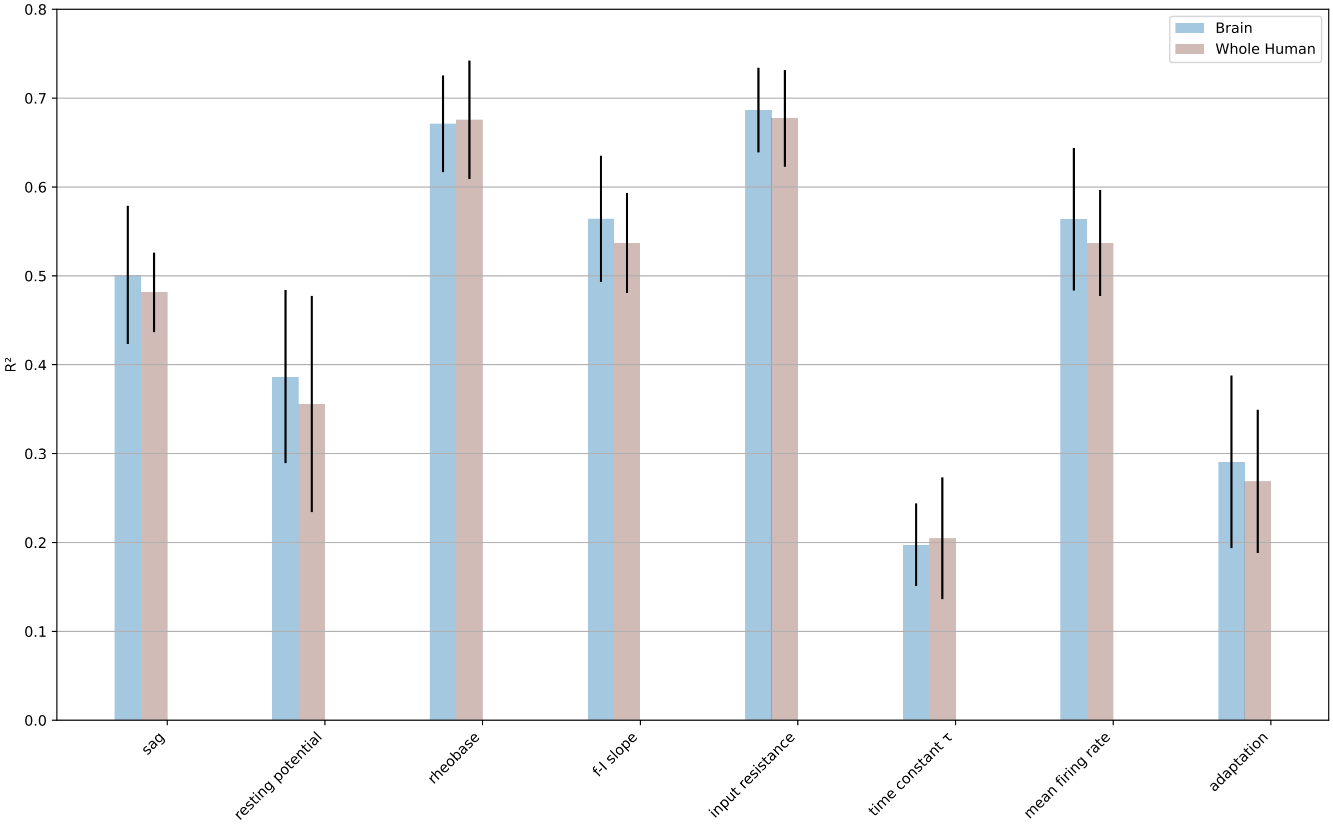


**Comparison of MLPs given exclusively scGPT embedding input: one using embeddings given by the whole human model and one using embeddings given by the brain model.** Error bars are standard deviations across the four folds of cross-validation.
